# Fragmentation and entanglement limit vimentin intermediate filament assembly

**DOI:** 10.1101/2022.03.19.484978

**Authors:** Quang D. Tran, Valerio Sorichetti, Gerard Pehau-Arnaudet, Martin Lenz, Cécile Leduc

## Abstract

Networks of intermediate filaments (IFs) need to constantly reorganize to fulfil their functions at different locations within the cell. The mechanism of IF assembly is well described and involves filament end-to-end annealing. By contrast, the mechanisms involved in IF disassembly are far less understood. *In vitro*, IFs are assumed to be very stable and their disassembly negligible. IF fragmentation has been observed in many cell types, but it has been suggested to be associated with active processes such as IF post-translational modifications. In this article, we uncover the contribution of filament spontaneous fragmentation in the assembly dynamics of type III vimentin IF using a combination of *in vitro* reconstitution probed by fluorescence imaging and theoretical modeling. We first show that vimentin assembly at low concentrations results in an equilibrium between filament annealing and fragmentation at times ≥24 hours. At higher concentrations, entanglements kinetically trap the system out of equilibrium, and we show that this trapping is reversible upon dilution. Taking into account both fragmentation and entanglement, we estimate that the mean bond breaking time is ∼18 hours. This translates into a mean breaking time of ∼ 5 hours for a 1 μm long filament, which is a relevant time scale for IF reorganization in live cells. Finally, we provide direct evidence through dual-color imaging that filament fragmentation and annealing coexist during assembly. By showing that IF fragmentation can occur without cofactors or post-translational modifications, our study provides new insights into the physical understanding of the IF length regulation.

## I. INTRODUCTION

Intermediate filaments (IFs), actin filaments and microtubules are the three main components of the cytoskeleton, which controls the mechanical properties and integrity of living cells [1, 2]. Cytoplasmic IF networks extend from the cell nucleus to the cell periphery and act in coordination with other types of cytoskeletal filaments to perform common cell functions such as cell migration, division and mechanosensing [3–9]. To perform these functions at different locations inside the cell, IFs need to constantly reorganize and remodel their networks. The main drivers of IF dynamics are active transport along F-actin and microtubules [10–16] and a combination of assembly and disassembly [14, 17–20]. In the case of vimentin, a type III IF, which is the major cytoskeletal component of mesenchymal cells [21], the mechanism responsible for filament assembly has been shown to involve longitudinal end-to-end annealing of filaments [14, 18, 22]. By contrast, the mechanisms involved in vimentin disassembly are much less understood [23]. Depolymerization events have been shown to be negligible in comparison with filament fragmentation [14], which is also referred to as severing in the literature [14, 17–19]. Vimentin fragmentation has been observed in many cell types [14], but it is unclear whether it occurs spontaneously or whether it requires other cues like cofactors or post-translational modifications [20, 24].

*In vitro* reconstitution experiments using recombinant vimentin have been instrumental in describing vimentin assembly, allowing to distinguish three steps: (i) after addition of salt which initiates the assembly, the lateral association of approximately 8 tetramers leads to the formation of 60 nm long unit-length filaments (ULFs) within seconds, (ii) ULFs anneal end-to-end to form extended filaments, (iii) these filaments undergo radial compaction, also referred to as maturation, within 10 to 30 minutes thus giving rise to 10-nm diameter filaments [25–32]. Mature filaments then further elongate by end-to-end annealing [30]. Spontaneous fragmentation has not been previously reported for wild type vimentin in purified systems reconstituted *in vitro* [31]. Based on this experimental evidence, theoretical models of vimentin assembly have consistently neglected the possibility of filament disassembly [28, 32–36]. They thus predict unlimited filament assembly without a finite steady-state length, in contrast to other cytoskeletal filaments. Available data, however, suggests that these assumptions may start to break down at a long time (over 4 h), where the quantitative accuracy of these models deteriorates [34, 35]. Data describing these long time scales is sparse, however, which contributes to our lack of understanding of filament disassembly.

Here, we investigate the reversibility of vimentin assembly using *in vitro* reconstitution and fluorescence microscopy coupled to theoretical modeling. We first study the assembly of vimentin IFs over long time scales, and observe that filament length gradually reaches saturation. This can be explained quantitatively by taking into account filament disassembly by fragmentation and, at high concentration, entanglement effects in the theoretical description. We then provide direct evidence that the assembly process is reversible by demonstrating a progressive decrease of the filament length after dilution. Finally, we show that filament disassembly is triggered by the fragmentation of individual filaments, which occurs concomitantly with end-to-end annealing during the assembly process. Combination of theoretical modeling and experiments allowed us to provide an estimation of the bond breaking time between two ULFs of ∼18 hours, which is of the same order of magnitude as the breaking time observed in cells [14].

## II. RESULTS

### A. Vimentin filament length reaches a plateau after ≥24 h assembly

To assess whether filament assembly can reach an equilibrium, we monitored the length of recombinant vimentin filaments over a period of more than 24 h after the initiation of assembly. Experiments were performed at 37 ^°^C in a sodium phosphate buffer (2.5 mM, pH 7.0), conditions in which the radial compaction is negligible (Fig. S1(a)). Previous works also showed that compaction is limited in this buffer condition and takes place in the first 10 minutes of assembly, which is very small compared to the time scale of our experiments [31, 37]. We used vimentin which was fluorescently labeled at the cysteine-328 (20 % labeling with AF488) to observe the filaments using fluorescence microscopy. We imaged the filaments at multiple time points and multiple concentrations from 0.01 mg · mL^−1^ to 1 mg · mL^−1^ (Fig. 1(a)) and quantified the mean length for each condition. The assembly curves showed that the assembly rate decreases over time and that the mean length reaches a plateau at times ≥24 h (Fig. 1(b)). For the highest concentration of 1 mg · mL^−1^, we could not provide length measurements after 6h assembly because the samples were too viscous to be aspirated in a pipette. We verified that the position of the fluorophore and the labeling rate had minor effect on the assembly dynamics, and especially the saturation length, using different fluorescence labeling methods (maleimide, which reacts with the cysteine-328 vs. succinimide ester, which mostly reacts with N-terminal NH2 groups) and different labeling rates (Fig. 1(c)). We further verified that filament assembly and length saturation were not impacted in the complete absence of fluorescence labeling using transmission electron microscopy (TEM) (Fig. 1(d)). Additionally, we confirmed that the sample mixing methods that we used in our experiments, *i*.*e*. pipetting and vortexing, did not affect the estimation of filament mean length (Fig. S1(b)). Finally, we verified that length saturation was not the consequence of vimentin proteolysis (Fig. S2). Thus, the existence of a plateau value for the mean length at long time scales suggests that filaments may have reached either an equilibrium involving the simultaneous assembly and disassembly of freely diffusing filaments, or a non-equilibrium (quasi-) steady-state where filaments are so entangled that their diffusion is severely hampered, hindering both assembly and disassembly [38].

**FIG. 1.**
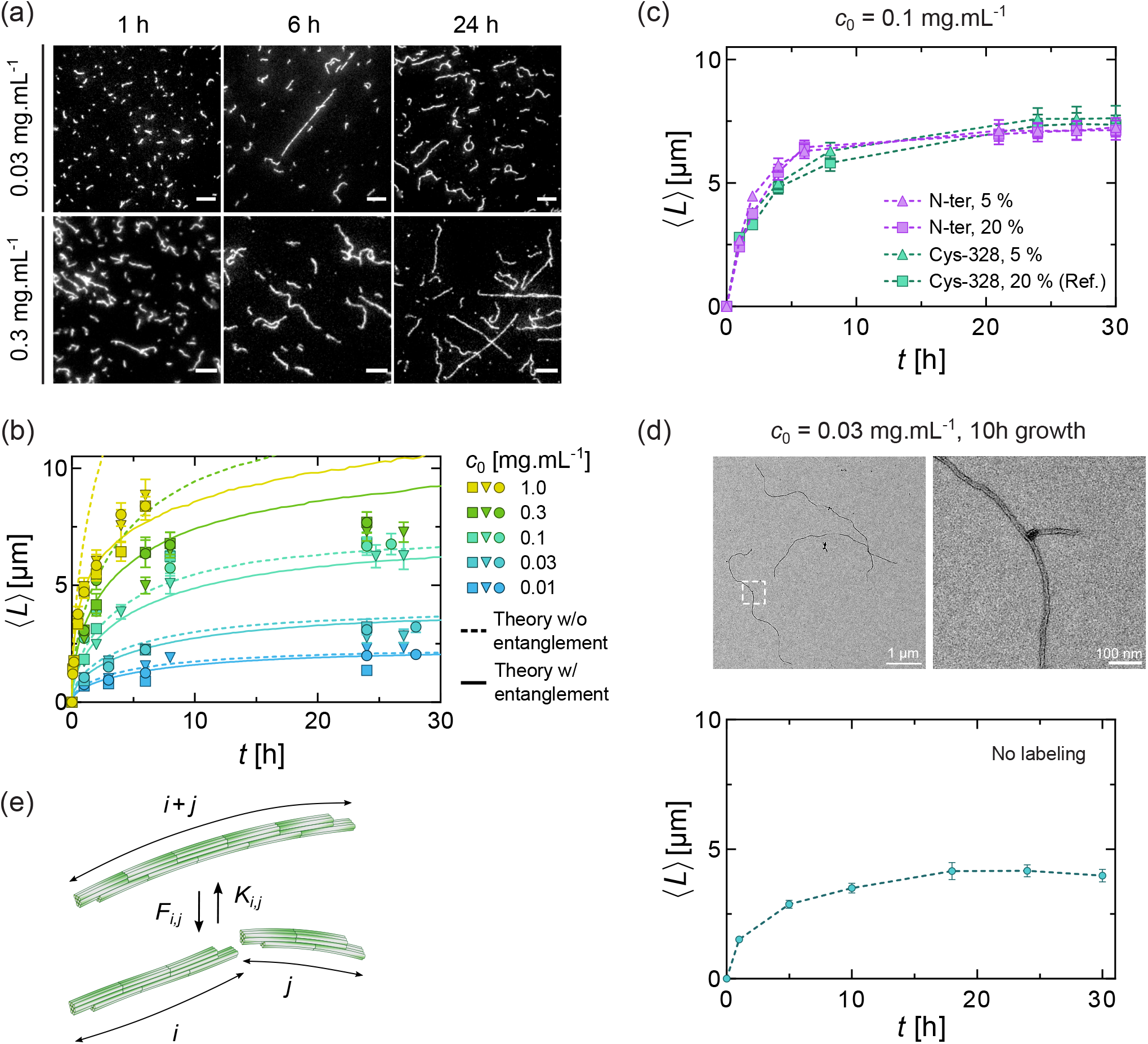
Filament length reaches a plateau during vimentin assembly. (a) Fluorescence images of *in vitro* vimentin filaments polymerized in assembly buffer at different initial concentrations, for a duration up to 28 h at 37 °C. For 0.03 mg · mL^−1^, filaments were fixed by mixing them with an equal volume of Glutaraldehyde 0.5 %. For 0.3 mg · mL^−1^, filaments were first diluted 20 times in assembly buffer before being fixed. Scale bar: 5 μm. (b) Graph of the mean length ⟨ *L* ⟩ of vimentin filaments at different initial concentrations *c*_0_ over time. The dashed lines correspond to theoretical fits from the model without entanglement, and the solid lines from the model with entanglement. Both fits were obtained *via* numerical simulations. The condition assembled at 0.1 mg · mL^−1^ was used as a reference for (c). (c) Graph of the mean length ⟨*L*⟩ of filaments assembled at the initial concentration of 0.1 mg · mL^−1^ with different fluorescence labeling methods (succinimide ester wich mostly reacts vimentin N-terminal NH2 groups and maleimide which reacts with the cysteine-328) and labeling rates over time. (d) Transmission electron microscopy image of unlabeled vimentin filament assembled at initial concentration of 0.03 mg · mL^−1^ for duration of 10 h at 37 °C and zoom of the boxed region. Graph of the mean length of unlabeled vimentin filaments as a function of time. Each data point in (b), (c) and (d) represents the average over 200 filaments and error bars represent the standard error of the mean. (e) Schematics illustrating the annealing at rate *K*_*i,j*_ and fragmentation at rate *F*_*i,j*_ used in the theoretical model.

### B. Modeling filaments as diffusing rods undergoing annealing and fragmentation accounts for the time evolution of filament lengths

To model filament growth in the presence of simultaneous assembly and disassembly, we used a mathematical model where our system is represented as a set of filaments of different lengths that randomly anneal end-to-end to form longer filaments, or fragment into smaller filaments. Filaments tend to become longer over time, implying that their diffusion slows down over the course of an experiment. As we are interested in the long-time behavior, we thus model their assembly as diffusion-limited; this assumption is also supported by the experimental data, as discussed below. We employed a generalization of the Smoluchowski coagulation equation which takes into account fragmentation [39] (detailed in Appendix A):

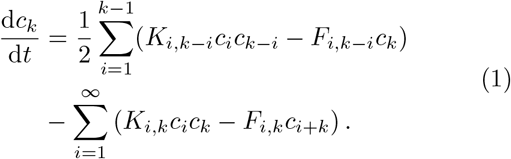

Here, *c*_*k*_ is the concentration of filaments of length *k* (in number of ULFs), *K*_*i,j*_ is the annealing rate between two filaments of respective lengths *i* and *j, F*_*i,j*_ is the rate at which a filament of length (*i* + *j*) fragments into two filaments of respective lengths *i* and *j*, and diffuse away (Fig. 1(e)). We note that events in which a filament is broken, but the fragments re-anneal before diffusing away, are not counted as fragmentation in this description (see Discussion). The dependence of the two rates on *i* and *j* encapsulates the key physical assumptions of our model. In a first version, we considered freely diffusing rigid filaments that associate end-to-end [40], yielding:

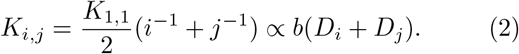

Here, *K*_1,1_ denotes the annealing rate for two isolated ULFs, and *b* is the ULF diameter. In general, *K*_1,1_ depends on temperature, on the viscosity of the buffer and on the size of the ULFs. We assumed the filaments obey Rouse dynamics, *i*.*e*., a simple model of free diffusion in the presence of a viscous friction against the surrounding fluid [41]. This yields a diffusion coefficient *D*_*i*_ ∝ *i*^−1^ for a filament of length *i* [41]. This set of assumptions implies that, in the absence of fragmentation, the mean filament length ⟨*L*(*t*)⟩ increases as *t*^1*/*2^ [42], consistent with our experimental data for time shorter than ∼10h when fragmentation is negligible (Appendix D). This sublinear increase of ⟨*L*(*t*) ⟩ implies a dependence of the reaction rate on filament length, supporting our assumption that the assembly is diffusion-limited. We note that the same model also gives a good description of the end-to-end annealing of actin [43]. In a passive system (e.g., in the absence of ATP hydrolysis), *F*_*i,j*_ is connected to *K*_*i,j*_ by the detailed balance requirement [39]:

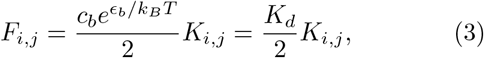

where *c*_*b*_ is a constant with the dimension of a concentration, *ϵ*_*b*_ *<* 0 is the energy change associated to the formation of a bond, *k*_*B*_ the Boltzmann constant and *T* the absolute temperature. In the second equality, *K*_*d*_ is the equilibrium dissociation constant. The mean length at equilibrium, ⟨*L*_∞_⟩, only depends on *K*_*d*_ and total concentration and not on the specific form of *K*_*i,j*_ [39]. We note that as a consequence of Eq. (3), the fragmentation rate depends on the size of the fragments. We solved Eq. (1) numerically using Direct Simulation Monte Carlo [44] (Appendix B). We fitted the theoretical predictions to the low-concentration experimental data by using two free parameters: the dissociation constant *K*_*d*_ and the mean bond breaking time (bond lifetime) 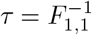. The fitting was performed as follows: For the three lowest concentrations (*c* = 0.01, 0.03 and 0.1 mg · mL^−1^) and for several *K*_*d*_ values, we performed 50 independent simulations, from which an average theoretical curve was obtained. We then fitted, for each one of the three experimental repeats, the theoretical curves to the experimental data. The fit was performed simultaneously over the three concentrations, so that for each experimental repeat we obtained, by minimizing the RMS residuals, a single best estimate of the dissociation constant and of the bond lifetime (see also Tab. I). Finally, the overall best estimates of *K*_*d*_ and *τ* were obtained by averaging over the best estimates obtained for each experimental repeat. These values were used to produce the theoretical curves reported with dashed lines in Fig. 1(b). For more details on the fitting protocol, see Appendix C.

**TABLE I.**
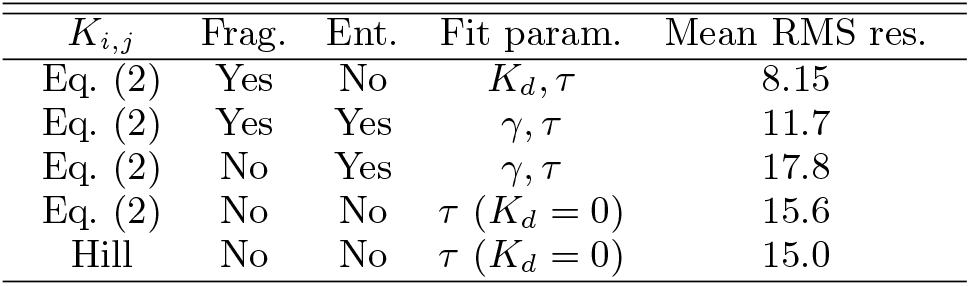
Mean RMS residual, averaged over the three experimental repeats, for the fits to the experimental data using different models. Column 1: *K*_*ij*_ used (our model, Eq. (2), or Hill model). Column 2: Fragmentation (yes/no). No fragmentation corresponds to *K*_*d*_ = 0. Column 3: Entanglement (yes/no). No entanglement corresponds to *γ* = 0. Column 4: Parameters used for the fit. Column 5: RMS residual from the fit averaged over the three experimental repeats. For more details on the fitting procedure, see Appendix C.

| $K_{i,j}$ | Frag. | Ent. | Fit param. | Mean RMS res. |
| --- | --- | --- | --- | --- |
| Eq. (2) | Yes | No | $K_d, \tau$ | 8.15 |
| Eq. (2) | Yes | Yes | $\gamma, \tau$ | 11.7 |
| Eq. (2) | No | Yes | $\gamma, \tau$ | 17.8 |
| Eq. (2) | No | No | $\tau$ ($K_d = 0$) | 15.6 |
| Hill | No | No | $\tau$ ($K_d = 0$) | 15.0 |

As shown in Fig. 1(b), dashed lines, for low concentrations (*c* = 0.01, 0.03 and 0.1 mg · mL^−1^), the model is in good agreement with the experimental data. The fitting procedure described above yields a dissociation constant *K*_*d*_ = (1.00 ±0.05) ×10^−5^ mg· mL^−1^ (mean ± SD, *N* = 3 repeats) equivalent to 6.1 ± 0.3 pM, and a mean bond breaking time 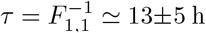. The resulting ULF annealing rate is *K*_1,1_ = 2(*τ K*_*d*_)^−1^ = (7 3) 10^6^ M^−1^ s^−1^. In Fig. 2(a) we compare the theoretical curves corresponding to *K*_*d*_ = 1.0 × 10^−3^ and *τ* = 13 h (solid lines), also shown in Fig. 1(b), to the curves obtained by setting *K*_*d*_ = 0, *i*.*e*., no fragmentation (dashed lines). For these, we find a worse quantitative agreement, *i*.*e*., a larger sum of square residuals (see Tab. I and also Appendix C). We also report (dotted lines) the comparison with the model of Hill [45] (here without fragmentation), which has been used in the literature [28, 32–34]. The comparison of this model and ours is detailed in Appendix E. We note that the Hill model gives an overall poorer description of the experimental data compared to ours.

**FIG. 2.**
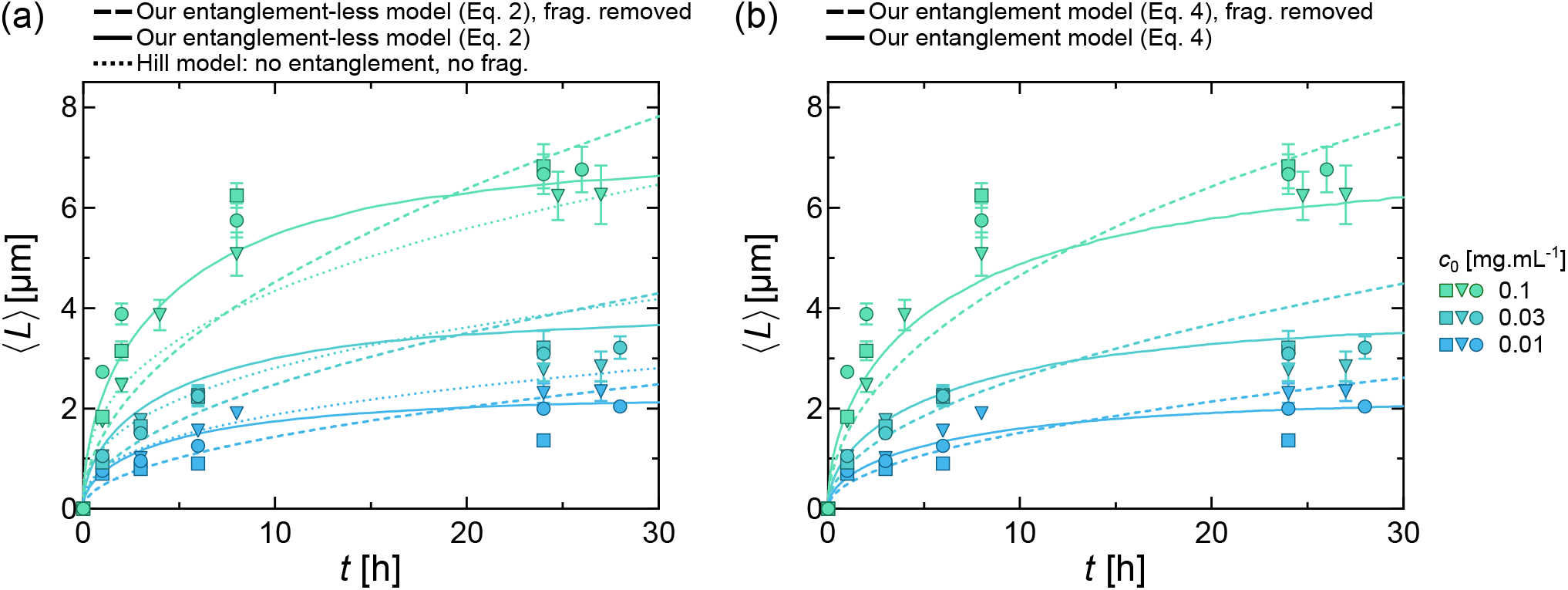
Comparison of different models at low concentrations (*c* = 1, 3, 10 equivalent to 0.01, 0.03 and 0.1 mg.mL^−1^ in experiments). **(a)** Models *without entanglement*. Dashed lines: Our model, *i*.*e*., *K*_*i,j*_ = *K*_1,1_(*i*^−1^ + *j*^−1^)*/*2, without fragmentation (*K*_*d*_ = 0). Solid lines: Our model with fragmentation; *K*_*d*_ = 1.0 × 10^−3^, *τ* = 13 h. Dotted lines: Hill model without fragmentation (*K*_*d*_ = 0). **(b)** Models *with entanglement*. Dashed lines: model without fragmentation (*K*_*d*_ = 0, *γ* = 7 × 10^−6^). Solid lines: model with fragmentation (*K*_*d*_ = 1.0 × 10^−3^, *γ* = 7 × 10^−6^). The values of *K*_*d*_, *τ* and *γ* used for the theoretical curves correspond to the average of the best fit estimates obtained by fitting each experimental repeat.

### C. Filament trapping by entanglement accounts for arrested assembly at high concentrations

At higher concentrations (*c* = 0.3 and 1.0 mg · mL^−1^), the filament lengths predicted by the model were consistently longer than those observed in the experiments. This led us to speculate that our assumptions of freely diffusing filaments break down at such high concentrations, and that filaments become increasingly entangled as they grow longer. This would lead to a kinetically arrested state, where steric interactions between the filaments prevent them from annealing. One of us has previously shown that the resulting slowdown in association rates due to the kinetic trapping of entangled rigid rods is well described by a simple analytical approximation [46]. A similar approach is relevant in our experiment, where at high concentration the persistence length of the filaments exceeds the mesh size of the system. To test our hypothesis, we modified our model to incorporate this mean-field kinetic trapping term in addition to fragmentation. The rationale of the model, is that two filaments that come into contact at a finite angle must align in order to achieve their annealing. The annealing is blocked if the ambient concentration of filaments is so high that other filaments stand in the way of this alignment, leading to a smaller, density-dependent reaction rate 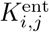. Here we summarize the derivation of 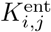, which is detailed in Appendix D.

To model the blocking resulting from entanglements, we assumed that two filaments can only anneal if the angle *θ >* 0 between them is smaller than some value *γ*. The reaction will be blocked if any other filament intersects the circular sector of area *A* = (*bl*_min_)^2^*θ/*2, with *bl*_min_ the minimum length of the two reacting filaments (see Fig. 7 in Appendix D). The probability that any given filament of length *k* intersects this circular sector, thus blocking the reaction, is *Abk/V*, with *V* the system’s volume. By averaging over all possible angles *θ*, it is possible to obtain an *average blocking probability*, ⟨*P*_*b*_⟩ = 1 − *g*(*x, γ*), where 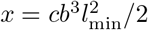, with 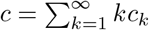 (total ULF concentration). Finally, the annealing rate for this model was obtained by multiplying *K*_*i,j*_ by 1 − ⟨*P*_*b*_ ⟩= *g*(*x, γ*), yielding:

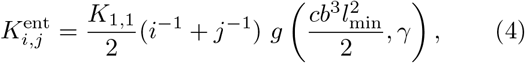

The function *g*(*x, γ*) = [1−(cos *γ* + *x* sin *γ*] *e*^−*γx*^)*/*((1+ *x*^2^)(1 − cos *γ*)) smoothly interpolates between 1 and 0 as *x* increases from 0 to ∞. This leads to an annealing rate identical to the one of Eq. (2) for low *c* and *l*_min_, and accounts for the slow-down of inter-filament reactions when these quantities are large. The fragmentation rate is still given by Eq. (3), with *K*_*i,j*_ replaced by 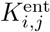, as is appropriate in the absence of ATP hydrolysis. We moreover kept the previously estimated value *K*_*d*_ = 1.0 × 10^−5^ mg · mL^−1^, as kinetic trapping should not affect the equilibrium properties of the system. A common fit over the five concentrations (*c* = 0.01, 0.03, 0.10, 0.30 and 1.00 mg · mL^−1^) averaged over the three experimental repeats, yields *γ* = (7 ± 2) ×10^−6^ rad. This small value for the angle *γ* suggests that filaments must be locally perfectly aligned in order for annealing to occur. As shown by the solid curves in Fig. 1(b), this improved model yields a significantly better agreement with the experimental data, with effectively only three free parameters being used to fit the five curves. We note that all the theoretical curves reach a plateau at long enough times, *i*.*e*., the system will eventually equilibrate. The new best estimate for the mean bond lifetime for this model is *τ* = 18 ± 4 h (mean ± SD, *N* = 3)(ULF annealing rate *K*_1,1_ = 2(*τ K*_*d*_)^−1^ = (5 ± 2) ×10^6^ M^−1^s^−1^). Note that entanglement has little impact on assembly dynamics at low concentrations, indicating that entanglement alone cannot explain the apparent length saturation with time in these conditions, and one must take fragmentation into account. This is shown in Fig. 2b, were we compare the theoretical predictions of the model with entanglement with and without (*K*_*d*_ = 0) fragmentation. For a 1 μm long filament, this translates to a mean breaking time of 5 ± 1 h (mean ± SD), as one can obtain by considering that the mean fragmentation time for a filament of length *k* is 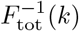, where 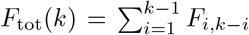 is the total fragmentation rate.

### D. The filament length distribution supports kinetic trapping at high concentrations

The two mechanisms for the saturation of the filament lengths imply differences in the filament lengths distribution in the long-time, plateau regime. At low concentration, the system reaches thermodynamic equilibrium. Assuming that the bonding free energy of two filaments does not depend on their lengths, this implies that the filament lengths are exponentially distributed [39]. At high concentration, small, highly mobile filaments react quickly, but the mobility and annealing rate of intermediate-length filaments is hindered by the surrounding tangle of other filaments. This situation implies a non-monotonic distribution of filament lengths, whereby short filaments are depleted while intermediate-length filaments accumulate due to their inability to react to form even longer filaments. To confirm this prediction, we investigated the distributions of filament length in the plateau regime (at *t* = 24 h) at low and high concentrations. Our observations show a good agreement with our model in both regimes, with the low concentration distribution being purely exponential (Fig. 3(a)) and the high concentration distribution showing a maximum (Fig. 3(b)), and thus confirm the existence of two distinct regimes for filament length saturation. Crucially, the maximum in the distribution at high concentration is only successfully reproduced when taking entanglement into account. It is due to the relatively faster depletion of short filaments when the reaction rate of the longer ones is heavily reduced by the entanglement, and provides a further indication that entanglements strongly influence the dynamic of polymer length in our system.

**FIG. 3.**
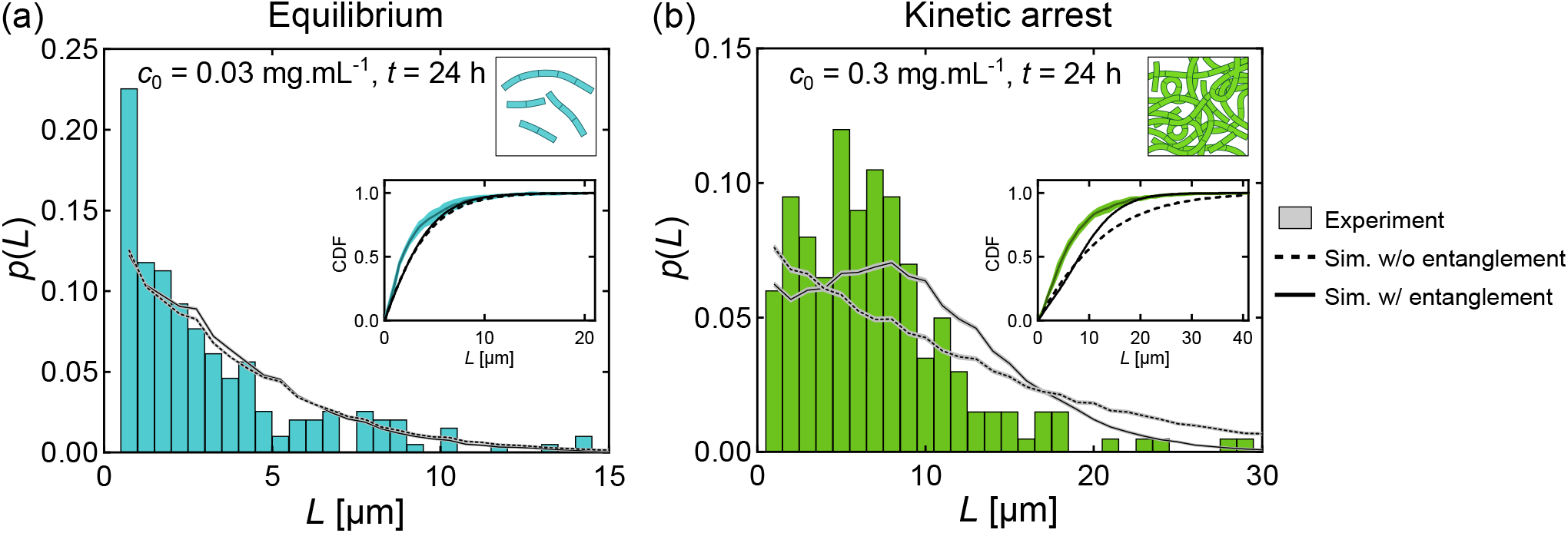
Length distributions after 24 h assembly. (a) Low concentration regime with initial concentration *c*_0_ = 0.03 mg · mL^−1^ and assembly duration of 24 h; (b) high concentration regime with initial concentration *c*_0_ = 0.3 mg · mL^−1^ and assembly duration of 24 h. Due to limitations in fluorescence imaging, the filaments with *L <* 0.5 μm are excluded from both experiments and theory. Sample size: ∼200 filaments. The theoretical curves have been obtained from two simulation models and averaged over 20 independent simulations. Dashed lines show the distribution when only considering annealing and fragmentation. Solid lines show the distribution with the addition of entanglement. Insets: cumulative distributions functions (CDF) of the filament lengths averaged over 3 experimental repeats at *c*_0_ = 0.03 mg · mL^−1^ (cyan) and *c*_0_ = 0.3 mg · mL^−1^ (green), averaged over 20 simulations without (dashed black lines) and with entanglement (solid black lines). The filled area around the lines depict the standard deviation over the 20 simulations or 3 experimental repeats. The experimental and simulated distributions were found to be non-significantly different at the 5% level using the Kolmogorov-Smirnov test (statistics *D* = 0.11 without and *D* = 0.10 with entanglement are above 0.05 for *c*_0_ = 0.03 mg · mL^−1^ and an experimental sample size of 675; *D* = 0.23 without and *D* = 0.21 with entanglement are above 0.04 for *c*_0_ = 0.3 mg · mL^−1^ and a sample size of 971).

### E. Vimentin IF assembly is reversible

The fact that filament length reaches an equilibrium at low concentrations implies that filament assembly should be balanced by disassembly. To clearly demonstrate that fragmentation is the mechanism responsible for disassembly, we shifted the equilibrium in the direction of filament shortening by diluting pre-assembled filaments. We diluted pre-assembled filaments at different dilution ratios and further incubated them at 37 °C for 6 h (Fig. 4(a)). We started from two different populations of filaments with a similar mean length of ∼3 μm, obtained either (i) after 2 h of growth with an initial concentration *c*_0_ = 0.2 mg · mL^−1^ where filament entanglement has limited effects on assembly or (ii) after 0.5 h of growth with a *c*_0_ = 1 mg · mL^−1^, when filament entanglement plays a role (Fig. 1(b)). Filaments shortened noticeably at low dilution ratio (1:10) and the shortening was even more pronounced at higher dilution ratio (1:100) in both conditions (Fig. 4(b)). Length quantification showed a gradual decrease of the vimentin IF length over time following dilution. Higher dilution ratios resulted in shorter filaments for the two populations of filaments, with low and high density of filaments (Fig. 4(c-d)).

**FIG. 4.**
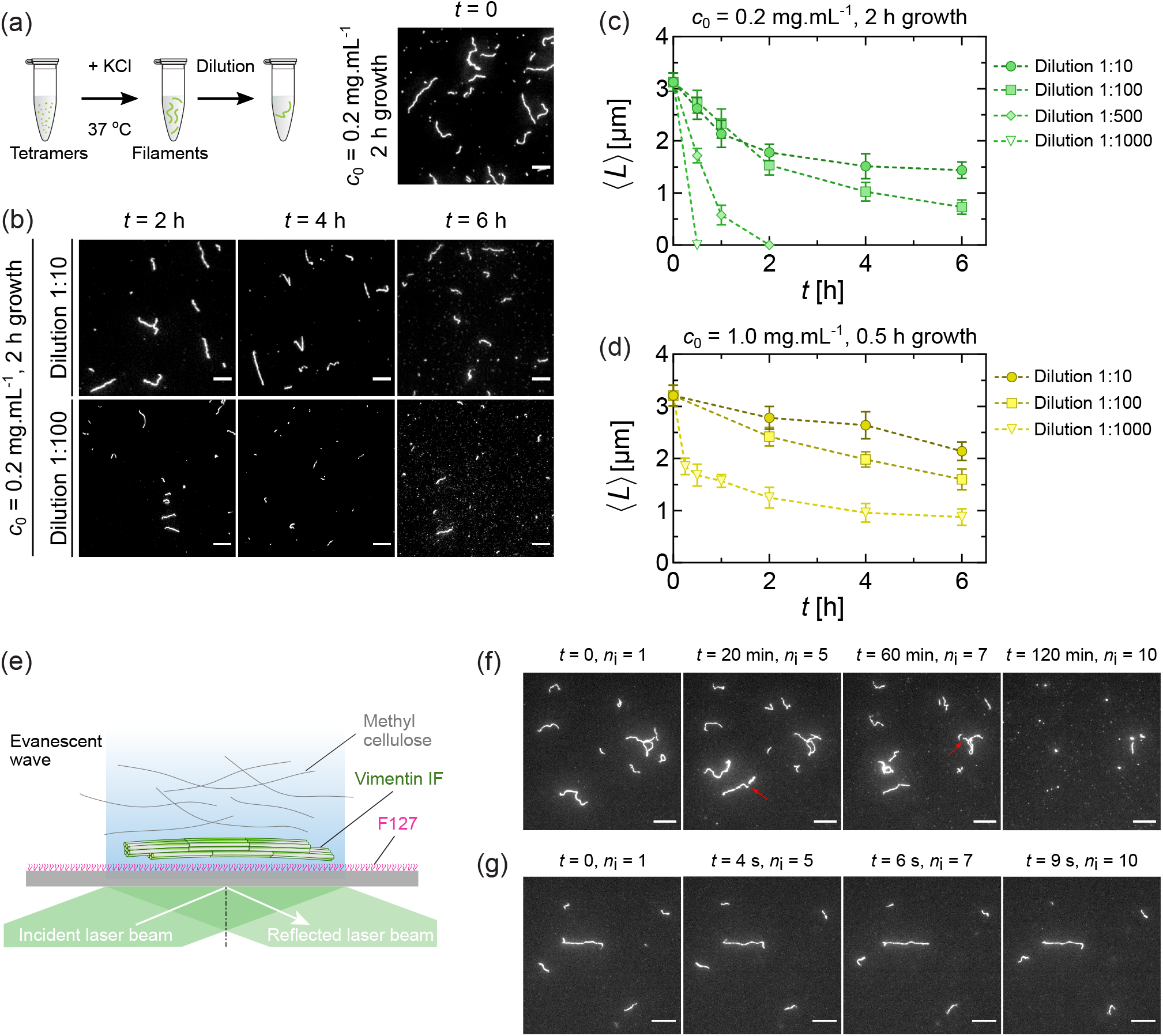
Vimentin assembly is reversible. (a) Schematics of the dilution experiment. Vimentin filaments were pre-assembled in the assembly buffer at 37 °C. The obtained filaments were then diluted at different ratios from 1:10 to 1:1000, and further incubated at 37 °C for up to 6 h. (b) Fluorescence images of vimentin filaments, with initial concentration 0.2 mg · mL^−1^ and assembled for 2 h, at different time points after dilution. Scale bar: 5 μm. (c) Evolution of mean length of diluted filaments with initial concentration 0.2 mg · mL^−1^ pre-assembled for 2 h, and (d) and 1.0 mg · mL^−1^ for 0.5 h. Error bars are standard deviations over 3 repeats. Sample size: ∼200 filaments per condition and repeat. (e) Schematics of the experimental setup comprising the flow chamber observed using TIRF microscopy. The glass coverslip is passivated by F127 preventing the non-specific adhesion of the filaments. Filaments are pushed close to the surface by the 0.2 % methylcellulose present in solution; they float without adhering to the surface. (f) Fluorescence images of diluted filaments in the flow chamber at different time points after dilution. Vimentin filaments were pre-assembled in the assembly buffer at 37 °C with initial concentration of *c*_0_ = 0.2 mg · mL^−1^ and 10 % labeling rate, then diluted at a ratio of 1:1000 in the assembly buffer with addition of 0.2 % methylcellulose and observed at ∼35 °C. Red arrows indicate the fragmentation events. *n*_i_ is the number of illumination times. (g) Phototoxicity control for the diluted filaments in the flow chamber. The filaments were imaged with a time interval of 1 s. We did not observe any filament breaking events after 10 illuminations. Scale bar: 10 μm.

To provide a direct observation of the fragmentation events in the dilution experiments, we designed an assay which allows us to observe the filaments *in situ* over time. We inserted the diluted filaments mixed with 0.2 % methylcellulose in a glass flow chamber whose surface was passivated by F127 in order to prevent filament attachment to the surface (Fig. 4(e)). By locating filaments close to the glass coverslips, methylcellulose allows their length to be followed over time. We observed fragmentation in time lapse movies of one frame every 20 minutes (Fig. 4(f), red arrows). We ruled out the possibility that the filaments fragmented due to phototoxicity by controling that continuous imaging did not induce filament breaking even after a large number of illuminations (Fig. 4(g)). Note that methylcellulose was used only for the experiments dedicated to the direct observation of fragmentation events in Fig. 4(e-g). Overall, these results demonstrate the reversibility of vimentin IF assembly both under conditions that lead to equilibrium and to the kinetically arrested state.

### F. Fragmentation and end-to-end annealing of vimentin IFs occur concomitantly

To obtain direct, filament-level evidence of the mechanisms at play during filament assembly, we performed dual color experiments that allowed us to follow the fate of segments of filaments as previously done in cells [14, 19]. We mixed two pre-assembled (3 h at 37 °C and 0.2 mg · mL^−1^) populations of vimentin filaments that were fluorescently labeled in different colors (green and red) and incubated them for another 6 h at 37 °C (Fig. 5(a)). Consistent with the observations of Fig. 1, we observed an increase of filament total length over time resulting from the annealing of filaments of different colors (Fig. 5(b-c)). We furthermore compared the length of single-color segments with the full length of the entire filaments and found that the mean length of the full filaments increases over time, the one of single-color segments decreases over time (Fig. 5(c)). These results show that vimentin filaments continuously fragment into shorter pieces while concomitantly re-annealing during the elongation phase. Moreover, we verified that fragmentation also takes place when filament length reaches saturation by mixing red and green filaments after 24 h assembly instead of 3 h (Fig. 5(d)). We observed a decrease of the single-color mean lengths, although less important than in (Fig. 5(c)), while the filament total length remains constant over time (Fig. 5(d)). These results indicate that filament entanglement obtained at long time scales limits both annealing and fragmentation at high concentrations as predicted by the theoretical model.

**FIG. 5.**
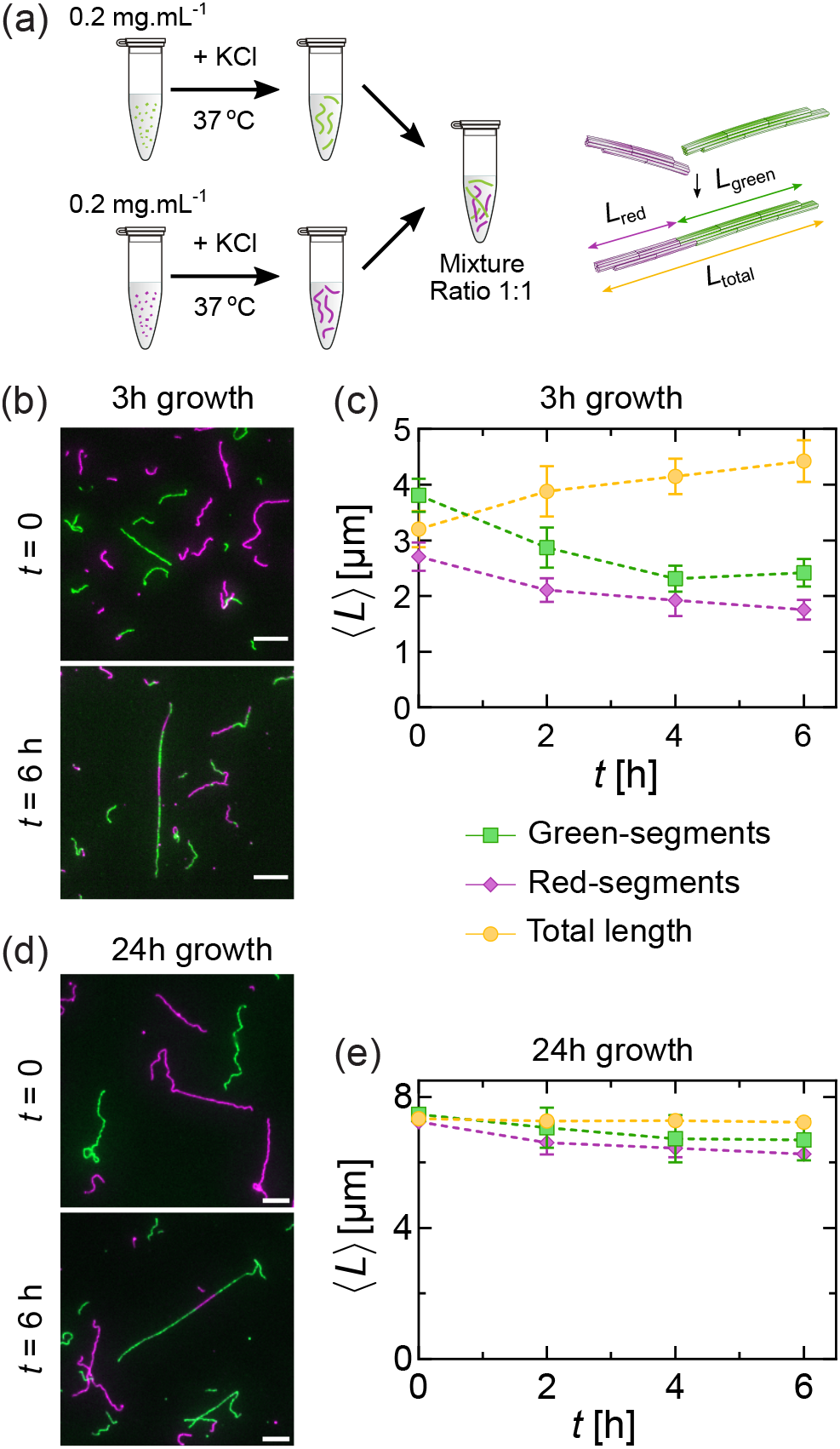
Annealing and fragmentation of vimentin filaments occur simultaneously during assembly. (a) Schematics of the dual color experiment. Two populations of vimentin filaments, both assembled from an initial concentration of 0.2 mg · mL^−1^, labeled in green and red, were pre-assembled at 37 °C for 3 h or 24 h. Then, they were mixed together at ratio 1:1 and incubated for another 6 h. (b, d) Fluorescence images at different time points after mixing and incubation for the filaments grown for 3 h (b) and 24 h (d). Scale bar: 5 μm. (c, e) Evolution of mean total length of filaments (yellow), length of red (magenta) and green segments (green) over time after mixing and incubation for the filaments grown for 3 h (c) and 24 h (e). Error bars are standard deviations over 3 repeats. Sample size: ∼200 filaments per condition and repeat.

## III. DISCUSSION

In this paper, we show that the vimentin filament assembly is limited by fragmentation and entanglement. We started by observing that the mean length of vimentin IFs saturated after 24 h of assembly. There could be several explanations for the filament length reaching a plateau: (i) the filaments become too long to diffuse and anneal over measurable time scales, (ii) the filament networks becomes too entangled, preventing annealing, or (iii) the filament disassemble, either by depolymerization from the extremities or by fragmentation, which would compensate the effect of annealing at long times. We tested these different mechanisms by theoretical modeling and accumulated experimental evidence to show that the only way to fully explain the experimental data is to take into account the disassembly by fragmentation. The reduction of the assembly rate due to the slower diffusion of long filaments, which is included in our model, was not sufficient to explain the length plateau without the inclusion of fragmentation (Fig. 2). Similarly, the effect of entanglement was found to be important at high concentration (*>* 0.1 mg · mL^−1^), but could not explain the plateau on its own without fragmentation, especially at low concentration (Fig. 2). Finally, the effect of depolymerization from filament extremities would decrease over time as the number of extremities decrease as a result of filament annealing, which was not compatible with the slowdown of assembly kinetics and the length saturation observed in the experimental data at long times.

Fitting all the experimental curves using theoretical modeling including fragmentation and entanglement allowed us to give an estimate of the bond breaking time between two ULFs of ∼18 hours. This time scale is of the same order of magnitude as the breaking time observed in cells [14]. Our results suggest that spontaneous fragmentation may play a role in intermediate filament reorganization within cells, independently from cofactors and post-translational modifications [20]. Phosphorylation events or proteins inducing fragmentation, like cofilin for actin filaments [47] or katanin for microtubules [48], may not be systematically involved in vimentin fragmentation in cells. Therefore, our results shine new light on the processes involved in vimentin turnover.

While the long time scales necessary to observe the impact of fragmentation are similar to those observed for vimentin fragmentation and annealing in cells, they are 6 times longer than the time scales explored in previous *in vitro* studies with comparable conditions (concentrations, buffer, assembly at 37 °C), which may account for the dearth of previous characterization of vimentin fragmentation [28, 34, 36]. Vimentin assembly has previously been probed over long time scales (*>* 144 h) in a study of the link between filament length and mechanical properties of the network. However, these experiments were carried out at room temperature, which strongly slows down the assembly dynamics [49]. It would be interesting to investigate the impact of temperature on filament fragmentation in more details in the light of our findings.

Our characterization of vimentin length saturation is reminiscent of observations on microtubules [50] and F-actin [51], although the lengths of the latter two reach a plateau more quickly (*<* 1 h) in comparable conditions of protein concentration, salinity and temperature. In addition, the mechanisms involved there are different and fundamentally active: dynamical instability in the case of microtubules and filament treadmilling in the case of F-actin, although annealing and fragmentation also play a non-negligible role for microtubules and actin as well [51]. Previous studies have shown that entanglements could induce kinetic arrest in dense actin networks in the presence of actin crosslinkers [38, 46]. In these studies, entanglements were shown to hinder the bundling of actin filaments. This bundling proceeds through a large-scale motion of filaments that become increasingly difficult in a dense network. The assembly of individual actin filaments was however largely unaffected by entanglement, due to the fact that it largely proceeds through the addition of single monomers, which can diffuse even in dense networks. By contrast, long vimentin filaments form through the annealing of shorter filaments, the diffusion of which can be severely hindered by entanglement. In the cancer cell line Hela, the concentration of intracellular vimentin has been estimated to be *>* 1 mg · mL^−1^ [52], but this level depends on the cell type, the differentiation state and the environment. Most of the vimentin proteins are assembled into filaments except a small fraction which can be found as soluble tetramers [53]. This suggests that filament entanglement might have an impact on vimentin assembly in the cytoplasm of cells with a dense network like Hela Cells. However, it may be overcome by other processes, such as active transport. Unfortunately, the high density of intracellular vimentin networks, the difficulty to follow filament tips in cells and resolution limitations of live microscopy make the quantification of filament length over time extremely challenging. Technological advances are therefore still needed to investigate the regulation of vimentin filament length in cells.

The spontaneous fragmentation of vimentin filaments may appear at odds with their well-characterized mechanical resilience, which allows them to undergo stretching by up to 300 % without breaking [54–58]. This apparent contradiction can however be resolved by noting the very different time scales involved in experiments where these behaviors are observed. Fragmentation is observed over several hours, while the mechanical properties of vimentin are typically probed over a few minutes at room temperature [54–58]. While the molecular mechanism responsible for vimentin longitudinal stretching has been shown to involve the unfolding of vimentin subunits [57, 59, 60], the molecular mechanisms responsible for filament fragmentation need to be characterized in more detail. One possible hypothesis builds on the observation that vimentin filaments can exchange subunits along their length during filament assembly [31], implying that subunits can spontaneously dissociate from the filament structure and re-associate with other binding sites. We speculate that these association/dissociation events could randomly result in the appearance of weak spots along the filaments, which could be responsible for filament fragmentation. Filament fragmentation may occur only if a large enough number of subunits are missing locally, which could account for its low rate of occurrence (timescale of tens of hours) if we take into account the fact that the subunit exchange rate is already very slow, *i*.*e*. ∼1 % exchange per hour [31]. This mechanism of fragmentation would also corroborate a previous report of vimentin filament polymorphism [61]. Measurements of the lifetime of tetramers within filaments could help validate this hypothesis at the molecular level. Moreover, since the phosphorylation of vimentin has been shown to regulate vimentin assembly by modifying the exchange rate of subunits towards a high off-rate in cells [62], it would be interesting to probe the impact of these post-translational modifications on the fragmentation mechanism. The softening of vimentin filaments observed after phosphorylation [63] may be also associated with an increased fragmentation rate. Other modifcations like S-gluathionylation at Cysteine-328 has already been shown to induce vimentin fragmentation [64].

The vimentin assembly dynamics observed in our experiments is well described by a mean-field model based on the Smoluchowski equation, which is based on the assumption that positions and orientations of the filaments are fully randomized within the network. In the model, the annealing rate of two filaments decreases when their lengths increase. This reflects the slower diffusion of long filaments compared to short ones, due to their larger friction with the surrounding fluid (see Eq. (2)). Based on this assumption and the detailed balance condition of Eq. (3), the total fragmentation rate 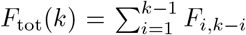 of a filament of length *k* does not increase linearly with *k*, as would be intuitively expected if all bonds broke independently and with the same rate. Instead, it increases much more slowly, specifically as ln(*k*), for long filaments. This behavior is however not incompatible with the afore-mentioned intuition. To understand this, we must note that the Smoluchowski formalism does not record microscopic bond-breaking events, but instead the instances where a bond breaks, then the resulting two filaments diffuse away from each other “to infinity”, *i*.*e*., by a length equal to several times their individual sizes. These latter instances are much rarer than the former in long, slowly-diffusing filaments. Indeed, most bond breaking events are followed by a re-binding event, rendered very likely by the resulting proximity of the filament ends. Likewise, the detailed balance condition (Eq. (2)) imposes that the kinetic slow-down factor g 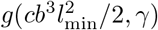 introduced in Eq. (4) applies equally to the fragmentation rate. Indeed, any physical process that hinders annealing must equally hinder the separation of the filaments following a bond breaking event, which increases the probability of their rapid re-binding. The success of this coarse mean-field approach to entanglements (Fig. 1(b)) suggests that the orientations of the filaments within the network remain largely random and isotropic throughout their assembly dynamics.

The annealing rate *K*_*i,j*_ used in our model is distinct from that of the Hill model [45], which is often used in the literature [28, 32–34] and posits *K*_*i,j*_ = ([ln(*i*) + 0.312]*/i* +[ln(*j*) + 0.312]*/j*)*/*(*i* + *j*) for rigid polymers undergoing end-to-end annealing. In particular, the Hill model predicts a slower filament growth than ours. To the best of our knowledge, none of the works that have made use of the Hill model to study the growth kinetics of vimentin have included fragmentation effects [28, 32–34]. We compared our model to a version of Hill’s with fragmentation and found that ours results in a better fit of the experimental data, especially in the early times *<*10h (Appendix E). This could be related to the implicit assumption in the Hill model that rotational and translational diffusion are decoupled. The Hill model without fragmentation displays an even worse agreement with the experimental data (Fig. 2). Our entanglement model with fragmentation improves the prediction of the experimental data compared to the Hill model, but the mean field approach we used does not entirely catch the kinetic slow down. A more refined model could be developed to describe more accurately the high concentration conditions, although this is beyond the scope of our current study..

Our study provides new evidence that vimentin filaments can fragment spontaneously, making the assembly process reversible. In the future, it would be of interest to provide a molecular understanding of filament breakage and probe how it can be regulated by post-translational modifications, which are known to impact filament disassembly. Moreover, the existence of fragmentation could be also tested in other types of IFs such as desmin or keratins to see to what extent it is a general feature. Finally, since more than 90 diseases have been associated with mutations of IF proteins and most impact the assembly process, it would be also interesting to probe specifically the effect of these mutations on the fragmentation rates. Overall, our results pave the way to new studies aiming at understanding the regulation of IF length, which plays a crucial role in IF dynamics and its related cellular functions.

## IV. MATERIALS AND METHODS

### *In vitro* reconstitution of vimentin wild-type

We purified vimentin wild-type protein from *E. coli* bacteria as described previously [65]. In short, vimentin wild-type was expressed in BL21 star (Sigma-Aldrich) cultured in Terrific Broth medium overnight at 37 °C, after induction at a DO of 1.2. We then centrifuged the culture medium to obtain the bacteria, then lysed them with lysozyme in the presence of DNase (Roche), RNase (Roche) and protein inhibitors (pefabloc and PMSF). We collected the inclusion bodies, washed them 5 times by successive steps of centrifugation and resuspension using a cooled douncer. After the last washing step, inclusion bodies were resuspended in a denaturing buffer (8 M urea, 5 mM Tris pH 7.5, 1 mM EDTA, 1 mM DTT, 1 % PMSF) and centrifuged at high speed (100 000 × g) for 1 h. After collecting the supernatant, we conducted vimentin purification after two steps of exchange chromatography, using first an anionic (DEAE Sepharose, GE Healthcare) then a cationic (CM Sepharose, GE Healthcare) column. The vimentin protein was collected in 2 mL Eppendorf tubes, and the concentration was monitored by Bradford. Only the most concentrated fractions were selected and mixed together. We stored the vimentin at -80 °C with additional 10 mM methylamine hydrochloride solution.

To obtain vimentin wild-type proteins for experiment, the denatured proteins were transferred to a dialysis tubing (Servapor, cut off at 12 kDa) and renatured by stepwise dialysis from 8 M, 6 M, 4 M, 2 M, 1 M, 0 M urea into sodium phosphate buffer (2.5 mM, pH 7.0, 1 mM DTT) with 15 minutes for each step. The final dialysis step was performed overnight at 4 °C with 2 L of the sodium phosphate buffer.

### Fluorescence labeling of wild-type vimentin

For the standard assembly assays, we labeled vimentin proteins by coupling their cysteine-328 with an AF-488 dye following a protocol that was described previously [30]. In details, denatured wild-type vimentin in 8 M urea was dialyzed for 3 h in a buffer containing sodium phosphate 50 mM, pH 7.0 and 5 M urea. Then, fluorescence dye (either AF-488 C5 maleimide or AF-555 C2 maleimide, ThermoFisher) dissolved in DMSO was added to the vimentin with a ratio between dye and vimentin molecule of 12:1, and mixed gently for 1 h at room temperature. The reaction was then quenched by addition of 1 mM DTT. The excess dye was removed from the mixture using a Dye Removal Column (#22858, ThermoFisher). Vimentin collected after the dye removal was renatured by stepwise dialysis from 8 M, 6 M, 4 M, 2 M, 1 M, 0 M urea to sodium phosphate buffer (2.5 mM, pH 7.0, 1 mM DTT). Renatured vimentin proteins were stored at 4 °C for up to a week. In our experiment, we noticed a small difference in assembly kinetics between two fluorescence dyes, AF-488 and AF-555, as shown in Fig. 4(c). To investigate the effect of the dye position on vimentin assembly kinetics, we used another labeling method with N-Hydroxylsuccinimide (NHS) ester. In a solution at pH 7.0, the succinimide dye compound forms a covalent bond mostly with the amino group at the N-terminal ends of the vimentin proteins. We followed the same labeling procedure except that we used AF 488-NHS ester instead of maleimide, and used a ratio between dye and vimentin molecule of 2:1.

### Assembly assays and sample fixation for imaging

For assembly of fluorescence-labeled vimentin filaments, non-labeled vimentin proteins were first mixed with AF-488-labeled proteins in sodium phosphate buffer (2.5 mM, pH 7.0) to the desired concentrations of 0.01, 0.03, 0.1, 0.3 and 1.0 mg mL^−1^ with a labeling rate of 20 %. The assembly was initiated by the addition of KCl at the final concentration of 100 mM. Incubation was performed at 37 °C for up to 28 h using a thawing water bath (Julabo Corio C, Seelbach, Germany). During the assembly, samples were taken from assembly solutions, then fixed with an equal volume of glutaraldehyde 0.5 % in the assembly buffer (2.5 mM sodium phosphate, pH 7.0, 100 mM KCl). In particular, for the high concentration of 0.1, 0.3 and 1.0 mg mL^−1^, samples were first diluted in the assembly buffer at ratio 1:10, 1:20 and 1:100, respectively, then fixed with an equal volume of glutaraldehyde 0.5 % in the assembly buffer. Similar to labeled filaments, non-labeled filaments were assembled at 0.03 mg mL^−1^ at 37 °C for a duration up to 30 h in the assembly buffer. During the assembly, small samples were taken at different time points, then fixed with an equal volume of glutaraldehyde 0.5 % in the assembly buffer. The fixed samples were then ready for imaging. No methylcellulose was used in the assembly assays.

### Fluorescence microscopy imaging

We transferred 3 μL of fixed vimentin filaments labeled with AF-488 or AF-555 onto a glass slide and put on a coverslip. The samples were then observed using an inverted microscope (Nikon Eclipse T*i*) and imaged using an sCMOS camera (C11440-42U30, Hamamatsu Photonics, Japan).

### Dilution experiment

We used vimentin labeled with AF-488 and a labeling rate of 20 %. We conducted the assembly experiments with an initial concentration of 0.2 mg mL^−1^ for 2 h and 1.0 mg mL^−1^ for 30 minutes, at 37 °C. Assembled filaments were then diluted at different ratios, from 1:10 to 1:1000 in the assembly buffer. We continued to incubate the diluted sets of filaments at 37 °C for another 6 h. Samples for imaging were taken every 1-2 hours. No methylcellulose was used in the dilution assays.

### Direct observation of diluted filaments

We demonstrated the filament fragmentation during dilution by direct observation of single filaments in a flow chamber. The flow chamber was constructed from silanized coverslips following a previously described protocol [66]. In details, we incubated cleaned coverslips with dichlorodimethylsilane 0.05 % in trichloroethylene for 1 h at room temperature. The coverslips were sonicated 3 times in methanol, each for 15 min. We then heated two silanized coverslips sandwiching pieces of parafilms up to 50 °C. The melted parafilm binding to the two coverslips resulted in a flow chamber. We passivated the flow chamber by flowing Pluronics F127 1 % in the assembly buffer and incubating for 1 h. Then, we thoroughly rinsed off the F127 by flowing an extensive amount of the assembly buffer. Finally, we diluted pre-assembled vimentin filaments (10 % fluorescence labeling rate, initial concentration 0.2 mg mL^−1^, assembly duration 3 h) 1000 times in the assembly buffer with addition of 0.2 % methylcellulose and flowed them in the chamber. We sealed the chamber and imaged the filaments using TIRF microscopy with heated objective and heating stage to maintain the temperature of the chamber at ∼35 °C. We captured images of the filaments every 5 min for the first 20 min, then every 20 min for the rest of the experiment. We performed a phototoxicity control where filaments were imaged every 1 s with the same number of illumination times as in the main experiment.

### Dual color experiment

We prepared two separate sets of vimentin, both at the same concentration 0.2 mg mL^−1^, but labeled in two different colors: one with AF-488 (green) and the other one with AF-555 (red). Both have the same 20 % fluorescence labeling rate. We conducted the assembly of the two sets for 3 hours at 37 °C. Then, we mixed the two sets of filaments together (mixing ratio 1:1), and continued to incubate the mixture at 37 °C for another 6 h. Samples for imaging were taken every 2 h. No methylcellulose was used in the dual color assays.

### Transmission electron microscopy imaging

We pipetted 4 μL of each fixed sample onto a carbon coated grid primarily glow discharged and incubated it at room temperature (25 °C) for one minute. Then, we performed negative staining by injecting 2 % uranyl acetate in water to contrast the grids and continue to incubate for one minute. The grids were then air-dried and observed under 120 kV using a Tecnai microscope (Thermofisher) and imaged using a 4k×4k Eagle camera (Thermofisher).

### Filament length analysis

Length quantification of vimentin filaments imaged in fluorescence microscopy and electron microscopy were conducted manually using Fiji. Only filaments above 0.5 μm length were considered for analysis in the fluorescence microscopy experiments.

## ACKNOWLEDGMENTS

We thank Brendan Evano, Harald Herrmann, Antoine Jégou, Stéphanie Portet, Guillaume Romet-Lemonne, Emmanuel Trizac, Raphaël Voituriez and Hugo Wioland for fruitful discussions, as well as Jean-Michel Camadro and Guillaume Jannot for technical help. Cécile Leduc thanks Tatjana Wedig and Harald Herrmann for teaching her the techniques involved in vimentin purification, and the support of the EU-supported Cooperation in Science and Technology (COST) action NANONET – Nanomechanics of intermediate filament networks. This project was funded by ANR-16-CE13-0019 (CL) and ANR-21-CE11-0004 (CL and ML), Marie Curie Integration Grant PCIG12-GA-2012-334053, “Investissements d’Avenir” LabEx PALM ANR-10-LABX-0039-PALM, ANR-15-CE13-0004-03, ERC Starting Grant 677532 (ML) and the 80—PRIME program of Centre National de la Recherche Scientifique (CL and ML). QDT and CL acknowledge the LabEx “Who am I?” (ANR-11-LABX-0071) and the Université Paris Cité IdEx (ANR-18-IDEX-0001) funded by the French Government through its ‘Investments for the Future’ program.

## APPENDIX

### A. Smoluchowski theory of annealing and fragmentation

In this Section, we introduce the Smoluchowski formalism to describe the annealing and fragmentation of vimentin filaments, and give more details on the interpretation of the Eqs. (1-3). The Smoluchowski equation allows us to describe a system of objects –here polymeric filaments– which undergo aggregation and fragmentation reactions. Each filament is characterized by its length (or, equivalently, mass) *k*, measured in ULFs, and the number density of filaments of length *k* is denoted *c*_*k*_. We assume that ULFs, *i*.*e*., filaments of length *k* = 1, cannot break apart. The annealing rate between two filaments of lengths *i* and *j* is denoted *K*_*i,j*_, whereas the fragmentation rate is denoted *F*_*i,j*_.

We make three main assumptions [39]: (i) No branching is allowed (each ULF can bind to two others at most), nor the formation of loops. (ii) The same free energy difference Δ*f*_*b*_ *<* 0 is associated with the formation of any bond. (iii) Detailed balance is respected, so that at equilibrium the rate of losing *k*-mers to fragmentation into *i*- and *j*-mers is exactly compensated by the rate of gaining *k*-mers from the annealing of *i*- and *j*-mers. Mathematically, the condition (iii) is expressed as follows :

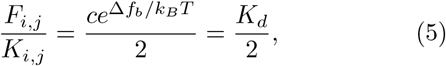

where *k*_*B*_ is Boltzmann’s constant, *T* is the absolute temperature, and *c* is the total number density of ULFs (similar to in Eq. (3). Here *K*_*d*_ is the equilibrium dissociation constant characterizing the reaction. We note that this quantity does not depend on the total ULF density *c*. To show this, we start by expressing *K*_*d*_, from Eq. (5), as:

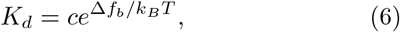

We can write this free energy change as the sum of an energetic term *ϵ*_*b*_ *<* 0 and an entropic term *T* Δ*s*_*b*_ *<* 0:

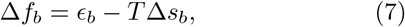

with

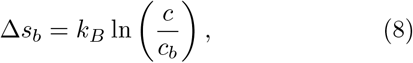

where 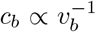, with *v*_*b*_ sometimes called the “bonding volume”[67]. We thus conclude that

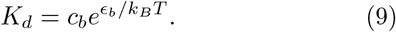

From the last expression, it is apparent that *K*_*d*_ only depends on the physics of the bonding process and on temperature, and not on the total density *c*.

Under the assumptions (i–iii) reported above, the concentrations *c*_*k*_(*t*) of filaments of length *k* are governed by the Smoluchowski equation Eq. (1) [39] where all the *c*_*k*_ depend implicitly on time and we have additionally assumed that *K*_*i,j*_ = *K*_*j,i*_ and *F*_*i,j*_ = *F*_*j,i*_. Here we consider a reaction rate which is appropriate for freely diffusing rigid filaments which undergo Rouse dynamics and anneal end-to-end. For this system, we expect *K*_*i,j*_ ∝ *b*(*D*_*i*_ + *D*_*j*_) [40], where *b* is the ULF diameter and *D*_*i*_ is the Rouse diffusion coefficient, which scales as *D*_*i*_ ∝ *i*^−1^ [68]. This amounts to applying the Smoluchowski formula, *K*_*i,j*_ ∝ (*D*_*i*_ + *D*_*j*_)*R*_*i,j*_, with *R*_*i,j*_ the target size (reaction radius) [69], to a target of size *b* diffusing with the diffusion coefficient of the whole filament. The modelling of the assembly process as diffusion-limited is justified experimentally by the fact that even at short times, the mean filament length increases sub-linearly with time (approximately as *t*^1*/*2^), implying that the reaction rate depends on filament length. For reaction-limited assembly, we would expect the reaction rate to be independent of filament length [69], and thus the mean length to increase linearly with time.

Following these assumptions, we have *K*_*i,j*_ = *K*_1,1_*/*2 (*i*^−1^ + *j*^−1^) (Eq. (2)). The dynamics is additionally constrained by the requirement that no ULFs are created or destroyed, *i*.*e*., the total mass of the system is conserved:

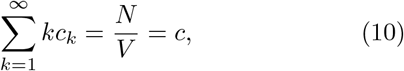

where *N* is the total number of ULFs and *V* is the system’s volume.

Eq. (1) implies that in a time interval Δ*t* and in the volume *V* we will have:

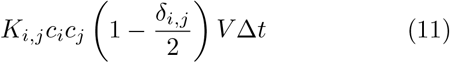

annealing events between an *i*-mer and a *j*-mer and

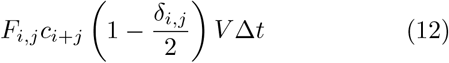

fragmentation events producing an *i*-mer and a *j*-mer, where *δ*_*i,j*_ is the Kronecker delta. The quantity in parentheses is introduced to avoid double-counting of reactions involving filaments of the same length (*i* = *j*). This is due to the fact that reactions with *i* = *j* are twice as rare as those with *i* ≠ *j*. This can be understood by considering, for example, a 4-mer and denoting the the probability per unit time that any of the three bonds in the 4-mer breaks as *p*_break_. One can then easily see that the probability per unit time that the 4-mer breaks into a 1-mer and a 3-mer is 2*p*_break_, whereas the probability that it breaks into two 2-mers is *p*_break_.

An important quantity we are interested in is the mean filament length at equilibrium, defined as

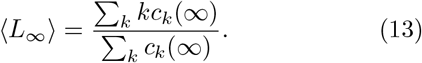

This quantity does not depend on the form of *K*_*i,j*_ (and *F*_*i,j*_), but only on the equilibrium constant *K*_*d*_ and on *c*, and it can be computed analytically. One finds [39]:

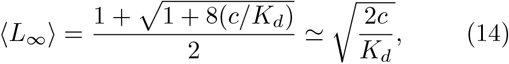

where the approximation is valid for large *c/K*_*d*_ (weak fragmentation). We will see in what follows that for the experimental system studied in this work, this approximation is valid.

In what follows, we will use for simplicity dimensionless quantities, defined by rescaling times by 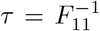 (the average breaking time of a bond between two ULFs) and all ULF densities by *c*_0_, the lowest ULF number density used in the experiments (corresponding to a vimentin concentration of 0.01 mg/mL). This yields, denoting the dimensionless quantities by a tilde:

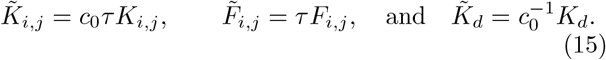

We note that with this choice of dimensionless quantities one has 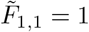 and thus

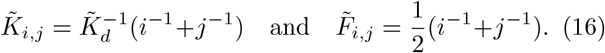

In the following Sections, we will drop the tilde when referring to dimensionless quantities for clarity.

### B. Description of the DSMC algorithm

Since the Smoluchowski equation Eq. (1) can only be solved analytically for certain forms of *K*_*i,j*_, we use here a numerical approach, called Direct Simulation Monte Carlo (DSMC) [44]. DSMC is a powerful stochastic method to solve differential equations such as Eq. (1), which samples the correct dynamics in the limit of large system sizes/large number of samples. We will here mainly follow the algorithm described in Ref. [44]: The starting point is an array **m** of length *N*, each element *α* of which contains a number *m*_*α*_ which represents the length of filament *α*:

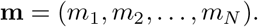

An element *m*_*α*_ = 0 represents the absence of a filament; moreover, we have 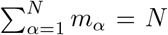 (total number of ULFs) because of mass conservation. Here, we choose *N* = 10^5^. To reflect the initial conditions in experiments, we additionally assume that the initial state of the system is

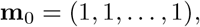

*i*.*e*., only monomers are present.

After the array **m** is initialized, we run the DSMC simulation, which consists in repeating a large number of times a Monte Carlo step (described below). The execution is arrested when the simulation time exceeds the equivalent experimental time.

With reference to Eq. (1), we define two quantities which will be useful in the description of the MC step below: 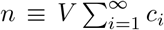 (total number of filaments) and 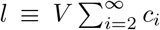 as (total number of filaments of length ≥2). We recall that only filaments of length 2 or more can undergo fragmentation.

Before the start of the simulation, we give an estimate of the maximum annealing rate *K*_max_ and of the maximum fragmentation rate *F*_max_. The exactness of the algorithm does not depend on this initial choice, however choosing values which are too far from the actual maximum rates can lead to a reduced efficiency [44]. For the annealing rate, Eq. (2), the maximum is by definition 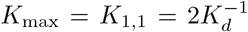 in dimensionless units. We thus have *F*_max_ = *F*_1,1_ = 1 as an estimate for the maximum fragmentation rate.

During every MC step, we either attempt to perform an annealing reaction (with probability *p*) or a fragmentation one (with probability 1 − *p*). The value of *p* is calculated initially and then updated during the course of the simulation in such a way that the average number densities *c*_*k*_ satisfy Eq. (1). At the beginning of each MC step, *p* is evaluated as

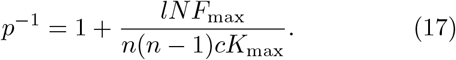

We will show below that this choice also guarantees that the simulation samples the correct number of fragmentation and annealing events per unit volume and unit time as are required by Eq. (1).

We define a *waiting time* variable that is set to zero at the beginning of the simulation. After each reaction, a waiting time increment is generated: These increments are also chosen in order to guarantee the correct number of annealing and fragmentation reaction per unit time/volume, as detailed below.

We can now describe the MC step, during which the following actions are performed:

1. We evaluate the probability of annealing *p* according to Eq. (17). The explicit form of *p*, Eq. (17), will be discussed in detail below.
2. We pick a random number 0 ≤ *R* ≤ 1 from a uniform distribution. If *R* ≤ *p*, we attempt a coalescence event:
  a. We pick a pair of elements of the array **m**, denoted *α, β* at random. Since there are *n*(*n* − 1) ordered pairs of elements to choose from, the probability to pick a specific pair is [*n*(*n* − 1)]^−1^. Let the length associated with these elements be *m*_*α*_ = *i* and *m*_*β*_ = *j*.
  b. We evaluate the annealing rate (in dimensionless units) as 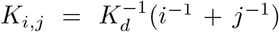. If *K*_*i,j*_ *> K*_max_, we set *K*_max_ = *K*_*i,j*_ and return to (1). Otherwise, we continue.
  c. We pick another random number 0 ≤ *R*^*′*^≤ 1 from a uniform distribution, and perform coalescence if *R*^*′*^≤ *K*_*i,j*_*/K*_max_. If coalescence is unsuccessful, we return to (1). Otherwise, we continue.
  d. We increment the waiting time by 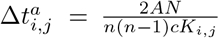. Here *A* is a parameter, the only condition on which is that it must be between 0 and 1, as we will discuss in more detail below.
  e. After updating the waiting time, we also update the array **m** by setting *m*_*α*_ = 0 and *m*_*β*_ = *i* + *j*.
3. If *R > p*, we attempt to perform a fragmentation event:
  a. We pick at random an element *γ* of the array **m**, with the condition that its length *m*_*γ*_ is equal to or larger than 2. The probability of choosing a particular element under this condition is *l*^−1^.
  b. We pick the length *i* of the first of the two fragments in which the filament will be fragmented (1 ≤ *i* ≤ *k* − 1) with probability (*m*_*γ*_ − 1)^−1^. The length of the second fragment is then *k*−*i*.
  c. We evaluate the fragmentation rate as *F*_*i,k*−*i*_ = [*i*^−1^ +(*k* − *i*)^−1^]*/*2. If (*k* − 1)*F*_*i,k*−*i*_ *> F*_max_, we set *F*_max_ = (*k* − 1)*F*_*i,k*−*i*_ and return to (1). Otherwise, we continue.
  d. We extract another random number 0 ≤ *R*^*′′*^≤ 1 from a uniform distribution, and perform fragmentation if *R*^*′′*^≤ (*k* − 1)*F*_*i,k*−*i*_*/F*_max_. If fragmentation is unsuccessful, we return to (1). Otherwise, we continue.
  e. We increment the waiting time by 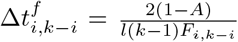, with *A* defined above in step (2). We will show below that choosing 1 − *A* here guarantees that the correct number of fragmentation events per unit time is obtained.
  f. After updating the waiting time, we also update the array **m** by setting the length of an element at random which has length 0 to *i* and setting *m*_*γ*_ to *k* − *i*.

Below, we prove that the definition of *p* (Eq. (17)) and the waiting time increments 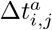 (for annealing) and 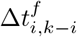 (for fragmentation) give a number of annealing and fragmentation events per unit time which is consistent with the Smoluchowski equation, Eq. (1).

Over a single MC step, the mean number of annealing events involving the pair (*α, β*) of elements of **m** with masses *m*_*α*_ = *i, m*_*β*_ = *j* is

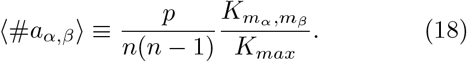

We note that in the algorithm we consider (*m*_*α*_, *m*_*β*_) as an ordered pair, and thus in Eq. (18) we consider the reaction (*i, j*) → *k* as distinct from (*j, i*) → *k*. The mean number of annealing events involving *any two filaments* with lengths *i, j* can be obtained by multiplying the above quantity by 2(1 − *δ*_*i,j*_*/*2)*V* ^2^*c*_*i*_*c*_*j*_. The factor 2(1 − *δ*_*i,j*_*/*2) takes into account the fact that, as mentioned above, for *I* ≠ *j*, there are two ways to perform the annealing, whereas for *i* = *j* there is only one. The factor *V* ^2^*c*_*i*_*c*_*j*_ is the product of the volume fractions of filaments of lengths *i* and *j*. We thus have

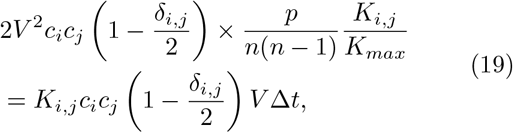

where we have equated the mean number of annealing events involving any two filaments with lengths *i, j* to the value required by the Smoluchowski equation, Eq. (11). From the equality Eq. (19) we obtain, recalling that *c* = *N/V*,

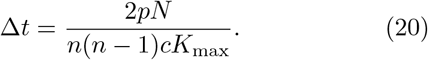

Eq. (20) relates the time interval Δ*t* to the probability of annealing. We will now obtain a second equality involving *p* and Δ*t*, which will allow us to prove that the expression Eq. (17) for *p* guarantees the correct number of fragmentation and coagulation events per unit time.

When performing a fragmentation event of a filament *γ* with mass *m*_*γ*_, we consider the two fragments of size *i, m*_*γ*_ − *i* in which it breaks as an ordered pair and thus *m*_*γ*_ → (*i, m*_*γ*_− *i*) is distinct from *m*_*γ*_ → (*m*_*γ*_− *i, i*). The mean number of fragmentation events involving filament *γ* where the first fragment of the ordered pair has length *i* is thus:

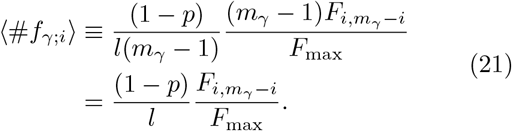

To obtain the mean number of fragmentations of a generic *k*−mer where any one of the fragments has length *i* we need to multiply this quantity by 2(1 −*δ*_*i,k*−*i*_)*V c*_*k*_, similarly to what we have done in the case of annealing:

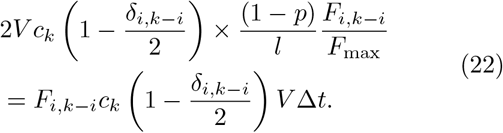

We have thus obtained a second equality involving Δ*t*:

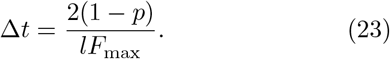

By equating the two expressions for Δ*t*, Eq. (20) and Eq. (23), we find Eq. (17). We have thus proven that the latter is the correct expression of *p*, which gives the correct number of fragmentation and annealing events per unit time and unit volume, as required by the Smoluchowski equation.

Finally, we will prove that the constants *A* and 1 − *A* introduced when calculating the waiting time increments are consistent with Eq. (20) and Eq. (23). To show this, it is sufficient to observe that the total time increment during a MC step is:

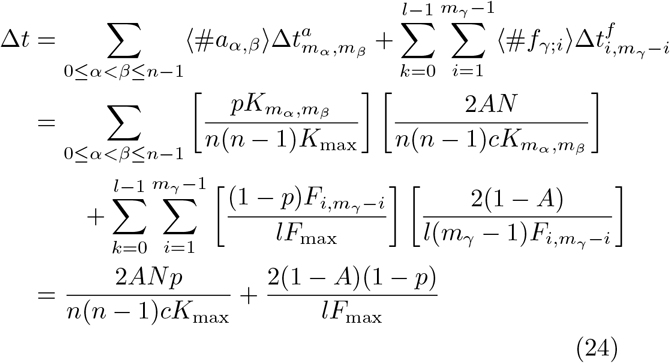

One can see that this equality is consistent with Eq. (20) and Eq. (23). We note that the algorithm samples on average the correct kinetics independently of the value of *A*, as long as 0 ≤ *A* ≤ 1. Here we take *A* = 1, meaning that the waiting time increment is calculated only after a successful annealing reaction, but not after a successful fragmentation reaction. This choice reduces the statistical noise on the data at short and intermediate times, where fragmentation is negligible and the filaments mostly undergo annealing reactions.

### C. Fitting the experimental data

In this Section, we describe the procedure followed to fit the experimental data shown in Fig. 1(b) of the main text (mean filament length *L* as a function of time) using the numerical results obtained with the DSMC simulations (Appendix B).

The objective of this fit is to determine the values of *K*_*d*_ (equilibrium dissociation constant) and *τ* (average bond breaking time) that best describe the experimental data. Since the data for *c >* 10 (corresponding to 0.1 mg/mL in experimental units) display the signature of a kinetic slowing down due to entanglement, as discussed in the main text, we have decided to fit only the data corresponding to the three lowest densities (*c* = 1, 3 and 10) to determine the values of *K*_*d*_ and *τ*. This is also justified *a posteriori* by comparing the fit results obtained by considering the three lowest, four lowest, or all the densities, as we discuss below.

We have run for each one of the densities *c* = 1, 3, 10 DSCM simulations for different values of the equilibrium dissociation constant *K*_*d*_. For each pair of (*c, K*_*d*_) values, in order to improve the statistics we have performed 50 independent simulations, in each one of which the random number generator was initialized with a different seed. From each simulation, we have obtained the mean length as a function of time, ⟨*L*(*t/τ*) ⟩ ; we have then averaged the ⟨*L*(*t/τ*) ⟩ curves produced from these simulations to obtain a single theoretical curve ⟨*L*_sim_(*c, K*_*d*_; *t/τ*) ⟩.

For each one of the three experimental repeats *R*_1_, *R*_2_, *R*_3_, we have then performed a common fit of the theoretical curves to the three experimental data sets corresponding to the densities *c* = 1, 3 and 10, in order to determine the best-fit value of *τ*. The experimentally measured mean filament length (in μm) has been converted to ULFs using the following relation, which takes into account the different effective length of a ULF when found along (49 nm) or at the extremity (59 nm) of a filament [60]:

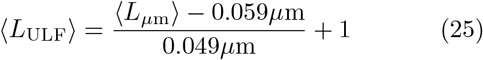

We note that, whereas fitting the three densities separately would result in three different estimates of *τ*, the common fit results in a single estimate of this parameter. The fit was performed using the non-linear least-squares method, implemented through the function *optimize*.*curve fit* of the open-source Python library Scipy [70], which employs the Levenberg–Marquardt algorithm [71]. We note that, in order to perform the fit, it has first been necessary to obtain continuous a representation of ⟨*L*_sim_ ⟩, so that its value could be calculated for an arbitrary *t/τ*. This was achieved by interpolating ⟨*L*_sim_⟩ with a cubic b-spline [72] (Scipy function *interpolate*.*splrep*).

The procedure described above has allowed us to find for each repeat and each *K*_*d*_ the value of *τ* which best fits the experimental data. We note that the fit has not been performed simultaneously on *K*_*d*_ and *τ* due to the fact that, whereas *τ* is a simple scaling factor of time, changing the value of *K*_*d*_ requires to perform a new simulation. In order to find the overall best-fit values of *K*_*d*_ and *τ*, for each value of *K*_*d*_ and for each repeat we have calculated explicitly the sum of the squared residuals [73], defined as:

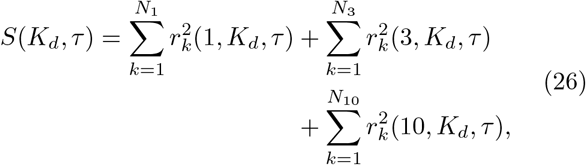

where

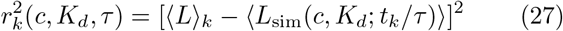

is the *k*-th squared residual, with (*t*_*k*_, ⟨*L*⟩ _*k*_), *k* = 1, …, *N*_*c*_ the experimental data points for a given density *c* and for the selected repeat. Each of the three functions *S*(*K*_*d*_, *τ*) corresponding to *R*_1_, *R*_2_ and *R*_3_ has then been minimized in order to determine the best-fit value of *K*_*d*_, denoted 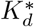. The corresponding *τ* value is also taken as the best-fit value of *τ* and denoted *τ*^*^.

In Fig. 6(a), we show for each of the repeats *R*_1−3_ the root-mean-squared (RMS) residual,

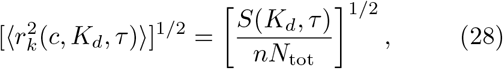

where *N*_tot_ = *N*_1_ + *N*_3_ + *N*_10_ and *n* is the number of densities considered (in the case of Eq. (26), *n* = 3). By direct inspection of the curves, we have found for 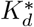 the values that are reported in Table II alongside the corresponding *τ*^*^ values. We have taken the overall best-fit value of *K*_*d*_ (*τ*) as the average over the 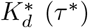 the final result is ***K***_***d***_ **= (1.00 *±* 0.05) *×* 10**^**−3**^ (mean ± SD). Converting this value to experimental units, one obtains ***K***_***d***_ **= (1.00 *±* 0.05) *×* 10**^**−5**^ **mg/mL**. The corresponding value of *τ* is ***τ* = 13 *±* 5 h**.

**TABLE II.** Best fit parameters 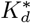 and *τ*\* from the common fit of the experimental data sets *c* = 1, 3, 10 for the three repeats *R*_1−3_ (see Fig. 6(a)).

| Repeat | $K_d^*$ | $\tau^*(\text{h})$ |
| --- | --- | --- |
| $R_1$ | $1.05 \times 10^{-3}$ | 8.32 |
| $R_2$ | $9.50 \times 10^{-4}$ | 14.3 |
| $R_3$ | $1.00 \times 10^{-3}$ | 17.3 |

**FIG. 6.**
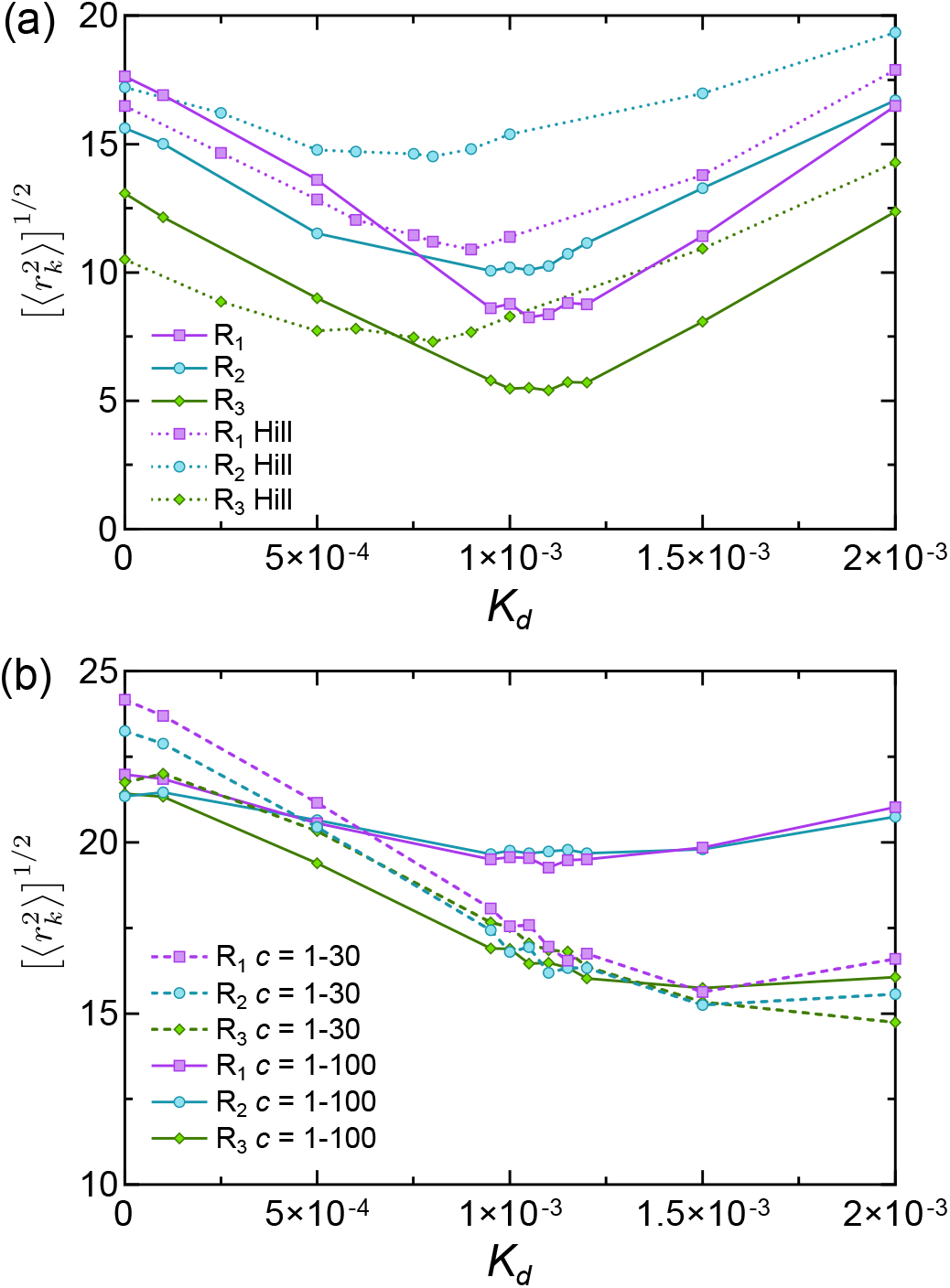
RMS residual, Eq. (28), as a function of the equilibrium dissociation rate *K*_*d*_. (a) Solid lines: Results from the common fit of the experimental data sets *c* = 1 − 10 for the three repeats *R*_1−3_. Dotted lines: Results for the Hill model with fragmentation, see Eq. (38) and Appendix E. (b) Results from the common fit of the experimental data sets for *c* = 1 − 30 (dashed lines) and *c* = 1 − 100 (solid lines) for the thre repeats *R*_1−3_.

The corresponding total fragmentation rate for a filament of *k* ULFs can be computed as

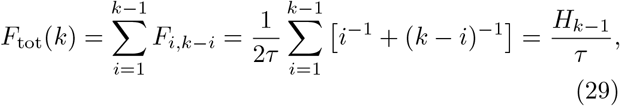

where *H*_*k*_ is the *k*-th harmonic number. For a filament of 1μm, corresponding to *k* ≃ 17 ULFs, for example, we obtain a mean fragmentation time 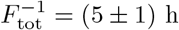.

In Fig. 6(b) we also show the functions 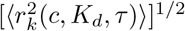 obtained by performing for each repeat a common fit including the density *c* = 30 (dashed lines) and one including *c* = 30 and 100 (solid lines). One can see that the values of *S* obtained from to these fits are overall significantly larger than those obtained by fitting the three lowest densities. This is due to the kinetic slowing down which affects the system for *c >* 10, as it was discussed in the main text (see also Appendix D), and justifies *a posteriori* the choice of determining *K*_*d*_ and *τ* from the three lowest densities only.

### D. Effect of entanglement (kinetic arrest)

Comparing with the experimental data the theoretical results obtained *via* the DSMC algorithm (dashed lines in Fig. 1(b) of the main text), one can see that the theory systematically and severely overestimates the mean filament length at equilibrium for the two highest densities (*c* = 30 and *c* = 100). We have speculated that this is due to the fact that at high density the experimental system becomes entangled, and thus the Smoluchowski description detailed in Appendix A becomes inadequate. Indeed, this theory is based on the assumption that annealing and fragmentation are purely two-body processes, which is a good approximation only in a dilute system. In a concentrated/entangled system, the presence of neighboring filament will likely hinder the annealing and fragmentation reactions. In the limit of very long filaments or very high concentrations, this will lead to a dramatic slowing down of the assembly, *i*.*e*. to a *kinetic arrest*. The objective of this Section is to propose a simple theoretical description of the microscopic mechanism of this slowing down. Since it would be very complex to extend the Smoluchowski theory to comprehend three-body interactions explicitly, we will adopt an “effective medium” description, in which the effect of the neighboring filaments on the annealing and fragmentation reaction is captured by a mean-field term.

Our model follows a similar approach to the one described in Ref. [46], which considers excluded volume interactions between rigid filaments undergoing bundling. In Ref. [46], all filaments are assumed to have the same length *L*, and to coalesce by lateral association. Here, we will assume the filaments to undergo end-to-end annealing and fragmentation as already described in Appendix A.

The microscopic mechanism that we propose for the hindering (*blocking*) of the microscopic annealing/fragmentation reactions is schematically represented in Fig. 7. In what follows, we assume that two filaments (modeled as diffusing rigid rods as detailed in Appendix A) can anneal/fragment only if the angle between them is smaller than *γ*, where *γ* is an adjustable parameter of the theory. We additionally assume an annealing/fragmentation reaction between two filaments of lengths (in ULFs) *i, j* will be blocked if at least another filament intersects the circular sector of area *A*. With reference to Fig. 7, one can see that

**FIG. 7.**
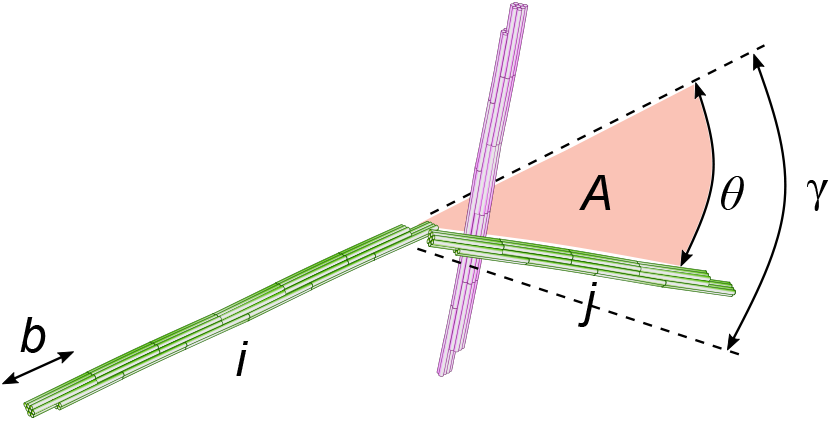
Schematic represenation of an annealing reaction between two filaments of *i* and *j* ULFs (green) being blocked by a third one (purple). The size of the ULF is *b*, and the angle *γ* represents the minimum angle required for the annealing to take place. If the third filaments lies in the shaded circular sector of area *A* = (*bj*)^2^*θ/*2, the reaction is blocked.

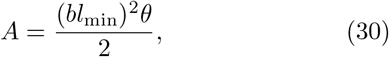

where *b* is the ULF size and *θ* the angle between the two reacting filaments, and we have defined *l*_min_ = min(*i, j*). The probability that a given filament of length *k* intersects this surface is

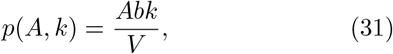

Thus, the probability that none of the *n* filaments in the system intersects the surface is, denoting the *blocking probability* with *P*_*b*_,

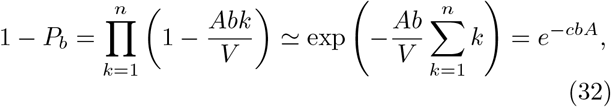

where *c* is the total concentration of ULF, and we have used the fact that *Abk/V* ≪ 1 for large volumes *V*. The probability of Eq. (32) was calculated for a given configuration of two filaments; thus, the average probability will be obtained by averaging over all the possible angles 0 *< θ < γ* between two filaments *i* and *j* undergoing annealing:

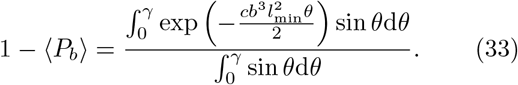

Solving the two integrals, we finally obtain

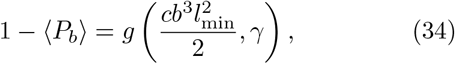

where

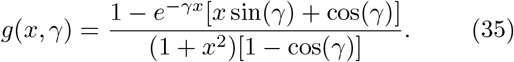

Thus, in conclusion, we propose that the annealing rate in the entangled regime is modified as follows:

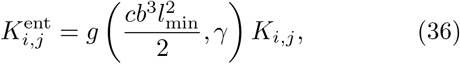

where *K*_*i,j*_ is given by Eq. (2). We note that lim_*x*→0_ *g*(*x, γ*) = 1, and thus in this limit we recover the model without entanglement.

To find the value of *γ* which best fits the experimental data, we have run DSMC simulations for different values of *γ*, while keeping the equilibrium dissociation constant the same as the one that was estimated by fitting the model without entanglement to the data for the three lowest densities. In particular, we have set *K*_*d*_ = 1.0 × 10^−3^, since this is the value which minimizes the sum of squared residuals when fitting the three lowest densities. The reason to keep *K*_*d*_ constant is that the kinetic slowing down caused by entanglement cannot modify the equilibrium properties of the system.

For each value of *γ*, we have run 50 independent simulations, in each one of which the random number generator was initialized with a different seed. We have then fitted the experimental data following the same procedure described above, considering this time the whole range of concentrations, *c* = 1, 3, 10, 30 and 100. When performing this fit, *K*_*d*_ has been treated as a constant, with *γ* and *τ* the fit parameters. As shown by the solid lines/filled symbols in Fig. 8(a), we have found that the best fit parameters, obtained by calculating the average and standard deviation over the three experimental repeats *R*_1−3_, are ***γ* = (7*±*2) *×* 10**^**−6**^ **rad** and ***τ* = 18*±*4 h**. From Eq. (29), the corresponding mean fragmentation time for a 1μm filament is 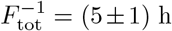. Since we had found *K*_*d*_ = (1.00 ± 0.05) ×10^−3^, we have also analyzed the effect that this 5% on *K*_*d*_ has on the estimates of *γ* and *τ*, finding that these do not change significantly (not shown).

**FIG. 8.**
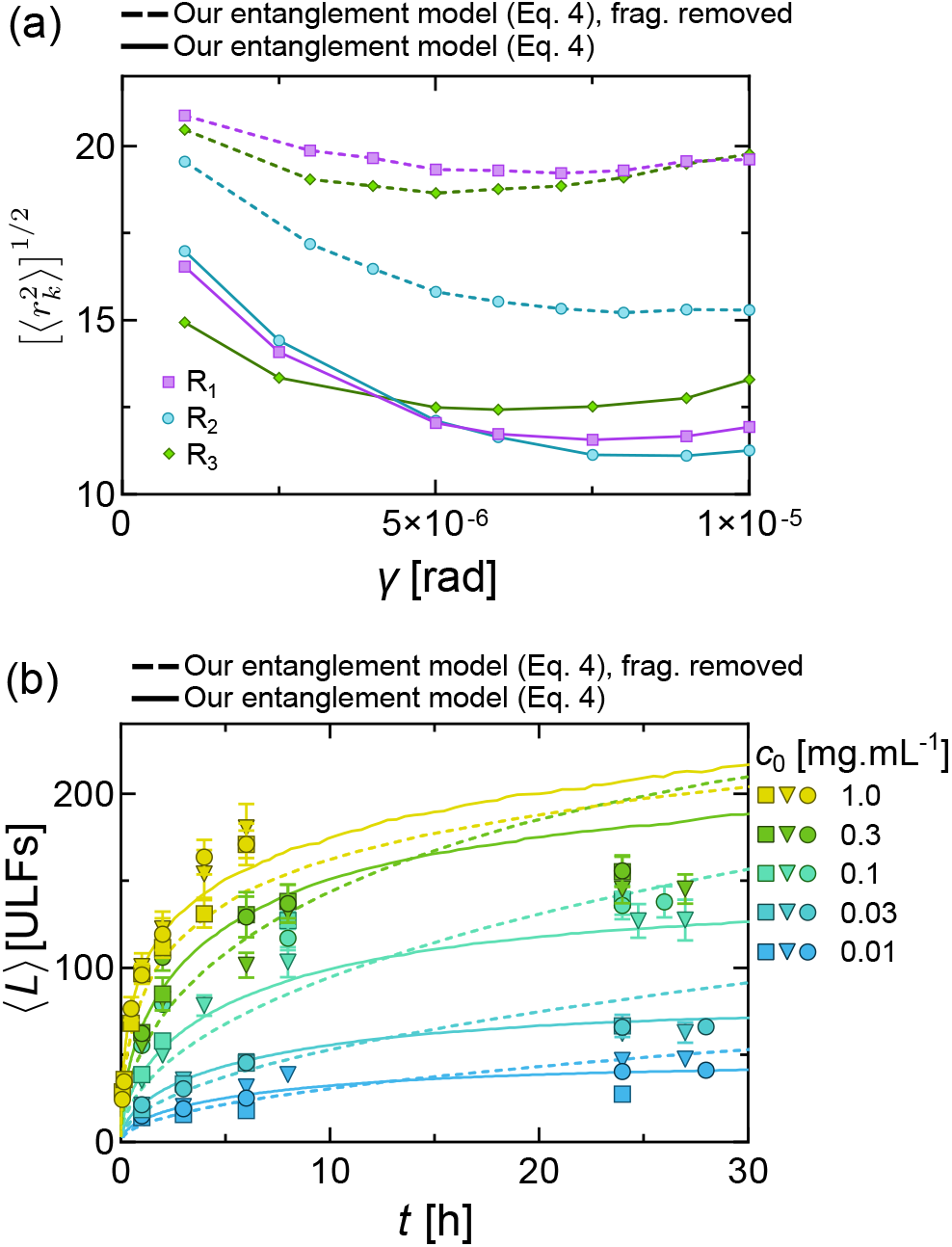
(a) RMS residual, Eq. (28), as a function of the minimum angle required for annealing *γ* (results from the common fit of the experimental data sets *c* = 1 − 10 for the three repeats *R*_1−3_). Solid lines: model with fragmentation and entanglement. Dashed lines: model without fragmentation, with entanglement. (b) Fit of the experimental data. Solid lines: model with fragmentation and entanglement (*K*_*d*_ = 1.0×10^−3^, *γ* = 7×10^−6^). Dashed lines: model without fragmentation, with entanglement (*K*_*d*_ = 0, *γ* = 7 × 10^−6^).

Finally, it is interesting to compare the experimental data with the same model as Eq. (36), but without fragmentation (*i*.*e*., *K*_*d*_ = 0). We have thus fitted the data with this model, finding, as shown by the dashed lines/empty symbols in Fig. 8(a), *γ* = (7 ± 2) ×10^−6^ rad. The value of *γ* for the model without fragmentation is thus identical (within the error bars) to the one found for the model with fragmentation. However, one can see that the RMS residual is significantly higher, and thus the model without fragmentation provides a worse description of the data. This is also shown in Fig. 8(b), where the fit results for the two models are compared. One can see that the model without fragmentation fails to capture especially the long-time saturation for *c <* 100, which is expected since fragmentation is more relevant at longer times.

### E. Comparison with the Hill model

In this Section, we will discuss how the experimental data compare to numerical results obtained using the Hill model [45] for the annealing rate *K*_*i,j*_. We find it especially relevant to discuss this model since it has been widely used for the analysis of vimentin growth kinetics [32–34]. We note that all of the previous studies have assumed the absence of fragmentation, whereas here we consider the Hill model *with fragmentation*.

The Hill model assumes that the two reacting filaments can be treated as diffusing rigid rods with diffusion coefficients *D*_*i*_, *D*_*j*_ and lengths *l*_*i*_, *l*_*j*_ = *b* · *i, b* · *j* (*b* = ULF size). Here *D*_*i*_ ∝ ln(*i*)*i*^−1^: although this relation describes somewhat more accurately the diffusion of thin rigid rods than the one used in this work [68], *i*.*e. D*_*i*_ ∝ *i*^−1^, the two have the same asymptotic behavior for large *i*. Moreover, the diffusion coefficient used by Hill vanishes for reactions involving single ULFs (*i* = 1 or *j* = 1), and thus in order to perform numerical calculations using this formula one must consider corrections to the logarithmic term [33, 34]. Here, following Portet *et al*. [34], we will use the following expression for *D*_*i*_ when discussing the Hill model:

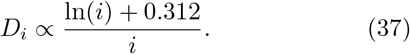

The constant 0.312 is a zero-order correction in *i*^−1^ for the diffusion coefficient of short filaments [74]. In the Hill model, two filaments that are at a distance *r*_*i,j*_ *<* (*l*_*i*_ + *l*_*j*_)*/*2 from each other will react with probability *p*_*i,j*_. This probability is evaluated by Hill as the fraction of surface of the sphere of radius *r*_*i,j*_ which is reactive. Since the area of the reactive site is ≈*b*^2^, one obtains 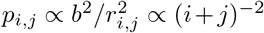. As we discuss below, this leads to a faster decrease of the reaction rate with filament length compared to our model. The reaction rate of Hill is thus:

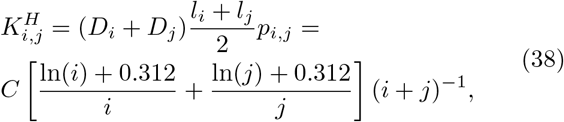

where *C* does not depend on *i, j* nor on density.

We have performed DSMC simulations of the Smoluchowski equation with fragmentation with the annealing rate given by Eq. (38). We recall that the fragmentation rate is derived from the detailed balance condition, Eq. (5). The simulations have been performed for the reduced densities *c* = 1, 3, 10, corresponding to the three lowest experimental densities, and for different values of the equilibrium dissociation constant *K*_*d*_. For each value of *c* and *K*_*d*_, we have run 50 independent simulations by initializing the random number generator with different seeds. In order to determine the best-fit value of *K*_*d*_ and the associated mean fragmentation time *τ*, we have followed the same procedure described in Appendix C. For the best fit values, we find ***K***_***d***_ **= (8.0*±*0.8) *·* 10**^**−4**^, corresponding to **(8.0*±*0.8) *·*10**^**−6**^ **mg/mL**, and ***τ* = 22*±*9 h**. The Hill model thus predicts a slightly smaller but comparable equilibrium dissociation constant and a larger mean bond breaking time than the one predicted by our model, Eq. (2).

In Fig. 6 we show the RMS residual from the fit, Eq. (28) (dotted lines, empty symbols), compared to the same quantity obtained with our model (solid lines, filled symbols). One can see by comparing the two curves that fitting the experimental data with the Hill model results in a larger value of the RMS residual in the range of *K*_*d*_ around the minimum 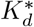.

The fit is shown in Fig. 9(a), where it is compared with the one obtained with our model. We note that the Hill model predicts a slower filament growth at short and intermediate times. This can be understood as follows: If one replaces for simplicity *i* and *j* with a single typical length *L* in Eq. (38), it is easy to see that for intermediate/long filaments thereaction rate scales as *K* ∝ *L*^−2^. In our model, Eq. (2), on the other hand, the scaling is *K* ∝ *L*^−1^. Thus, the Hill model predicts a faster decrease of the reaction rate with increasing filament length.

**FIG. 9.**
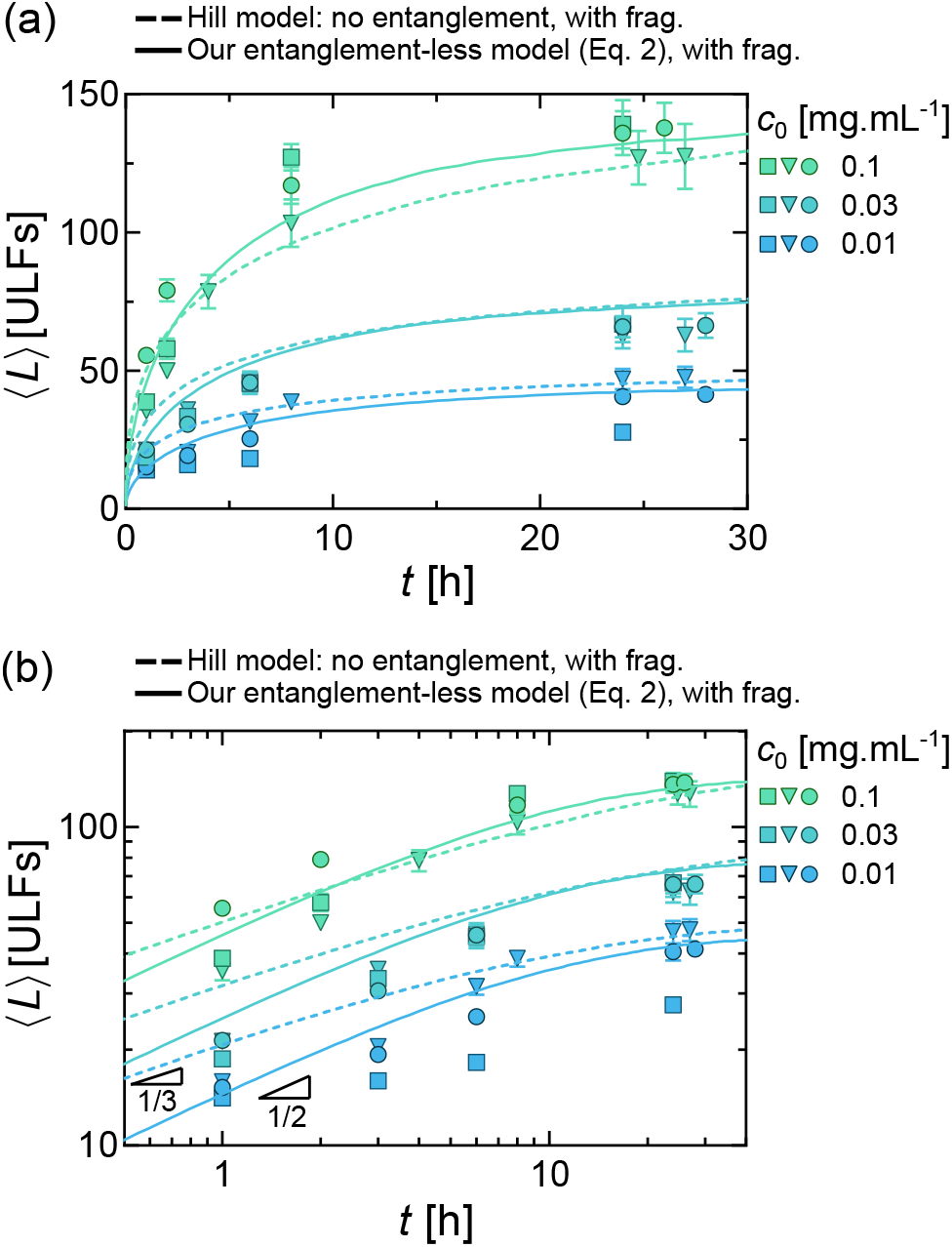
(a) Fit of the experimental data (symbols) obtained using our model for the annealing rate, *K*_*i,j*_ = *K*_1,1_(*i*^−1^ + *j*^−1^)*/*2 (solid lines), and the Hill model with fragmentation (Eq. (38), dashed lines). (b) Same data as in (a) in double-logarithmic scale. For our model, *K*_*d*_ = 1.00 ×10^−3^, *τ* = 13 h. For the Hill model, *K*_*d*_ = 8.0 × 10^−4^, *τ* = 22.0 h. Note that all the data shown in A-B are for models *without entanglement*, as the latter are not relevant in this range of densities.

It can actually be shown that, in the absence of fragmentation, a scaling *K* ∝ *L*^−*λ*^ leads to an increase of the mean filament length with time as *L*(*t*) *t*^1*/*(1+*λ*)^ [75]. This results in *L*(*t*) ∝ *t*^1*/*2^ for our model and *L*(*t*) ∝ *t*^1*/*3^ for the Hill model. These two regimes are indeed observed at times *<*10 h, when fragmentation is less relevant, as shown in Fig. 9(b), where we show the same data as in Fig. 9(a) in double-logarithmic scale. The experimental data follow a slope much closer to the value 1*/*2 predicted by our model than to the 1*/*3 predicted by the Hill model. We thus conclude that our model captures the assembly kinetics better than the Hill model.

## Supplementary figure

**Figure S1:**
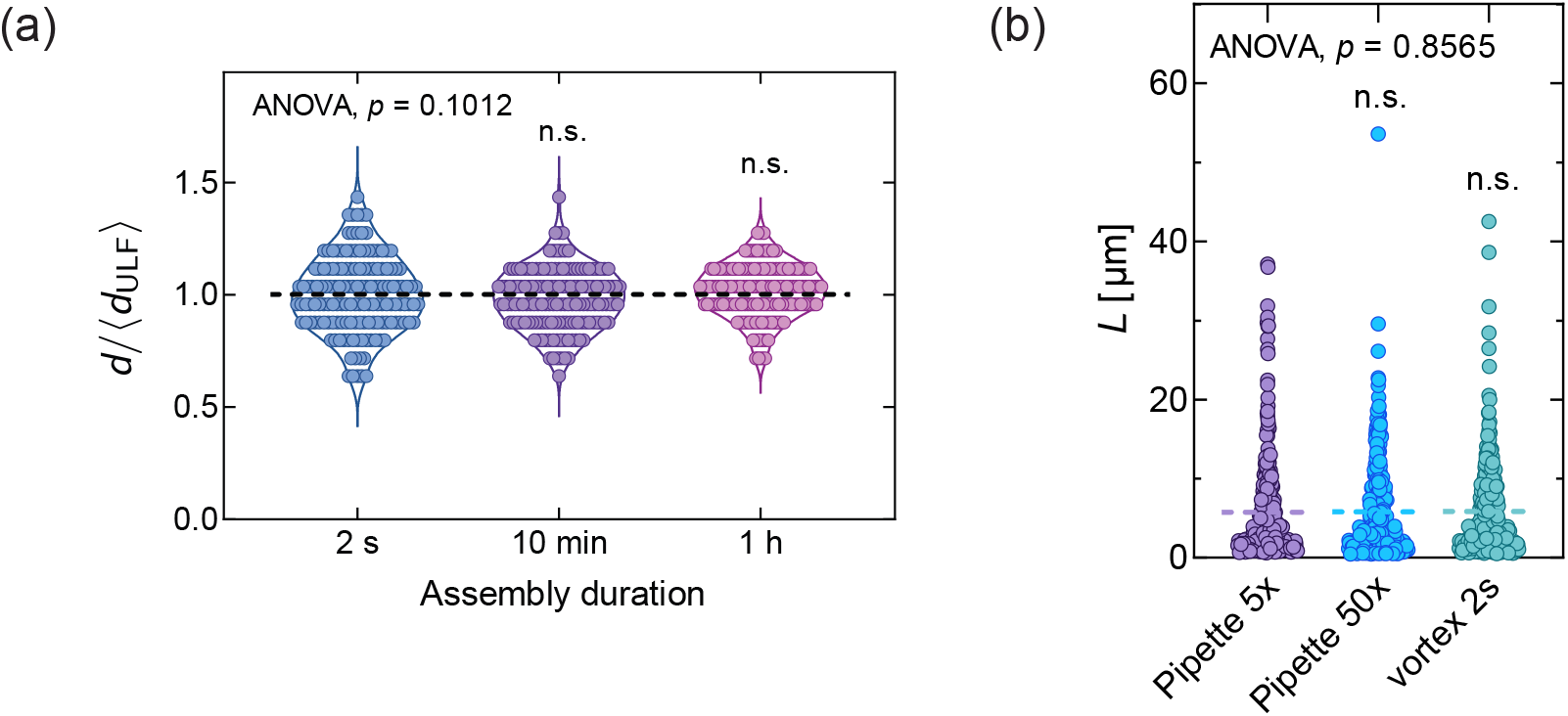
Quantification of filament diameter and controls for filament mixing. (a) Comparison of filament diameter at different assembly durations. Vimentin filaments were assembled at initial concentration of 0.1 mg · mL^−1^ at 37 °C and samples were collected at different assembly durations (2 s, 10 minutes and 1 h). Filament diameter was measured from transmission electron microscopy images acquired the same day with the same defocus, and normali:.ed to the average diameter of ULFs obtained at 2 s assembly, 12.5 nm. Sample si:.e: 150 filaments. (b) Comparison between 3 methods of mixing filaments during sample preparation. Vimentin filaments were pre-assembled with initial concentration of 0.3 mg·mL^−1^ for 4 h at 37 °C. Dashed lines correspond to the median value. Filament length was measured from fluorescence images of filaments diluted from the same batch by 3 different mixing methods. Sample size: > 300 filaments.

**Figure S2:**
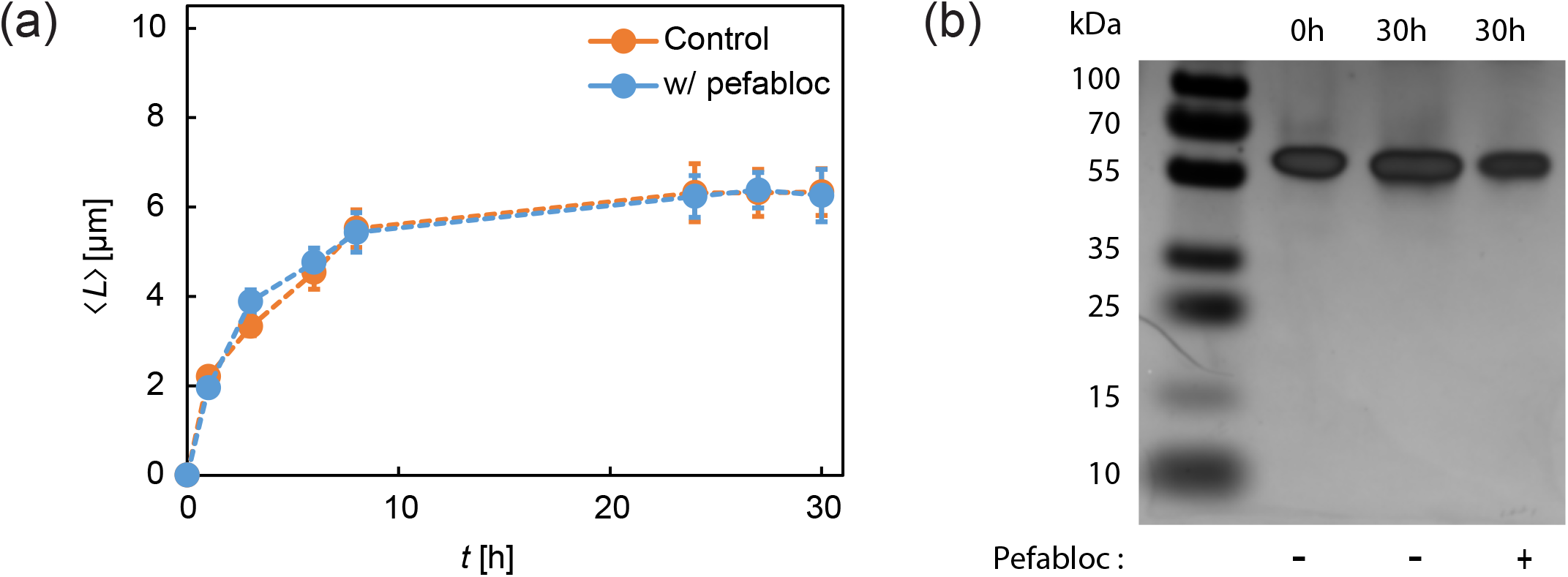
Proteolysis has no impact on the saturation of filament length. (a) Graph of the mean length ⟨*L*⟩ of filaments assembled at the initial concentration of 0.1 mg · mL^−1^ with and without 0.5 mM pefabloc, an inhibitor of serine proteases, as a funtion of time. Fresh pefabloc, which has a half-life time of 6h, was added every 6h. Sample size: > 200 filaments per condition. Error bars represent the standard error of the mean. (b) Silver stained SDS-page gel of vimentin before assembly, after 30h of assembly and after 30h assembly in the presence of 0.5 mM pefabloc. There is no evidence of degradation bands.

